# NLR DNA copy-number expansion outpaces accumulation of predicted coding-intact copies

**DOI:** 10.64898/2026.09.10.750634

**Authors:** Liming Xiong, Xuexiao Zou, Feng Liu

**Author notes:** Correspondence: Feng Liu XueXiao Zou.

## Abstract

Whether nucleotide-binding leucine-rich repeat (NLR) DNA copies and predicted coding-intact copies accumulate proportionally within homologous neighbourhoods remains unresolved. We compared these counts across 173 anchor-defined neighbourhoods in 11 *Capsicum* genomes. After accounting for neighbourhood and genome effects, predicted coding-intact copies increased less than proportionally with DNA copies. DNA-copy doubling corresponded to 1.48-fold and 1.84-fold increases in expected coding counts under strict and inclusive definitions, respectively. The strict association persisted across sequence and sampling controls, with highly expanded arrays and heterogeneous coding recovery affecting interpretation. An independently defined panel of 17 *Arabidopsis thaliana* accessions reproduced sublinear scaling using curated gene counts. Matched pepper RNA-seq showed a transcript-support gradient across predicted coding states. At DH06, a near-megabase segmental duplication in *C. pubescens* contained intact and disrupted family pairs. Copy-specific HiFi reads placed the same F396 coding lesions in both physical blocks. Together, these results reveal a recurrent relationship between NLR array size and coding composition in the surveyed plant systems. Interpreting DNA expansion together with coding state provides a more informative basis for comparing NLR repertoires and prioritizing immune-gene candidates.

## Introduction

Nucleotide-binding leucine-rich repeat (NLR) receptors are central components of plant intracellular immunity, but their evolutionary repertoires encompass more than the sequences that encode active receptors (Jones et al. 2016; Monteiro and Nishimura 2018). The birth-and-death framework describes resistance-gene families as products of duplication, divergence and loss (Michelmore and Meyers 1998). Foundational work in lettuce established that resistance-gene clusters can span several megabases and that unequal crossing-over and gene conversion can remodel their members (Meyers et al. 1998; Chin et al. 2001). Duplication therefore changes the amount and arrangement of receptor-associated DNA, whereas preservation of an intact coding sequence is a distinct outcome. An expanded array can retain intact copies, copies with explicit coding lesions and sequences whose coding status remains unresolved.

Pan-NLRome studies have made this diversity accessible beyond single reference genomes. Species-wide inventories in *Arabidopsis thaliana* and genome-assisted analyses in *Solanum americanum* have revealed extensive presence–absence and allelic variation (Van de Weyer et al. 2019; Lin et al. 2023). Within *Arabidopsis* clusters, expansion can be concentrated in a few radiating lineages rather than distributed evenly among all members, complicating reference-based orthology assignment (Lee and Chae 2020). Hypervariable receptor subfamilies also differ from more conserved NLRs in their sequence-diversity profiles (Prigozhin and Krasileva 2021), consistent with the heterogeneous evolutionary behaviour documented within lettuce resistance-gene clusters (Kuang et al. 2004). These findings make local genomic context and the identity of the counted sequence central to comparisons of NLR repertoires.

Recent studies have analysed coding degradation in its genomic context. Teasdale et al. (2025) annotated intact and degraded NLRs within 121 pangenomic neighbourhoods from 17 *Arabidopsis* accessions, showing that multiple metrics are needed to describe NLR diversity. Cacao comparisons likewise linked local duplication to extensive copy-number variation and pseudogene-rich arrays (Winters et al. 2025). At broader phylogenetic scales, microsynteny-informed classification and pennycress pangenome comparisons demonstrated that chromosomal position can retain information not captured by gene-level sequence similarity alone (Guo et al. 2025; Bird et al. 2026). These studies establish a positional view of NLR diversity that incorporates coding degradation, motivating an explicit analysis of how coding composition changes with DNA-copy number.

Specifically, within homologous neighbourhoods, do predicted coding-intact copies increase in direct proportion to DNA copies? Observing fewer coding copies than DNA copies does not distinguish proportional from sublinear accumulation: the same deficit occurs when a constant fraction of an expanding array remains coding-intact. A declining fraction instead implies less-than-proportional accumulation of coding-intact copies. Testing this distinction requires separately measured DNA and coding counts, a comparison unit that does not depend on intact target-gene annotations, and explicit treatment of positions that cannot be evaluated. The resulting comparison describes the coding composition of extant arrays.

The large, repeat-rich genomes of *Capsicum* provide a contrasting setting in which to examine this relationship. Comparative studies have documented dynamic NLR repertoires and a contribution of retroduplication to resistance-gene expansion in pepper (Seo et al. 2016; Kim et al. 2017; Kim et al. 2021). Chromosome-scale, graph-pangenome and telomere-to-telomere resources now span cultivated peppers and divergent relatives (Liu et al. 2023; Chen et al. 2024; Zhang et al. 2025). A recent T2T pan-NLRome catalogued 4,789 annotated genes in 227 orthogroups across 11 pepper genomes (Dong et al. 2026). These resources enable a complementary analysis of DNA-defined copies, including sequences without complete gene models. Complex tandem arrays remain difficult to annotate consistently (Lim et al. 2026). Annotation-independent NLR detection provides an entry point (Steuernagel et al. 2020), while coding recovery must be evaluated separately to avoid interpreting annotation failure as biological degradation.

Here, we test proportional coding-copy accumulation within conserved non-NLR anchor-defined neighbourhoods across 11 *Capsicum* genomes. Log-link count models distinguish a constant conditional coding fraction from sublinear scaling, while sequence, sampling and held-out recovery analyses assess how measurement and comparison design affect the result. The independently defined *Arabidopsis* neighbourhood panel tests recurrence in a second plant system. Matched transcript data characterize the RNA support associated with each genomic coding state. Finally, local alignments and copy-specific long reads resolve intact and disrupted family pairs within a near-megabase duplication at DH06. These analyses connect comparative count scaling with the coding composition of physical copies.

## Results

### Conserved chromosomal positions reveal NLR copy and coding-state diversity

Homologous NLR neighbourhoods differ in DNA copy number and coding composition. We measured positional homology, DNA presence and predicted coding state separately using conserved non-NLR flanks (CaNLOG, *Capsicum* NLR Orthologous Genomic-neighbourhood framework). In one observed Chr03 neighbourhood, Grif1614 and Zhangshugang carry two and four DNA-NLR copies, respectively, yet each retains one strict coding-intact call; incomplete anchor support leaves the corresponding Andean position non-evaluable (Fig. 1a).

**Figure 1.**
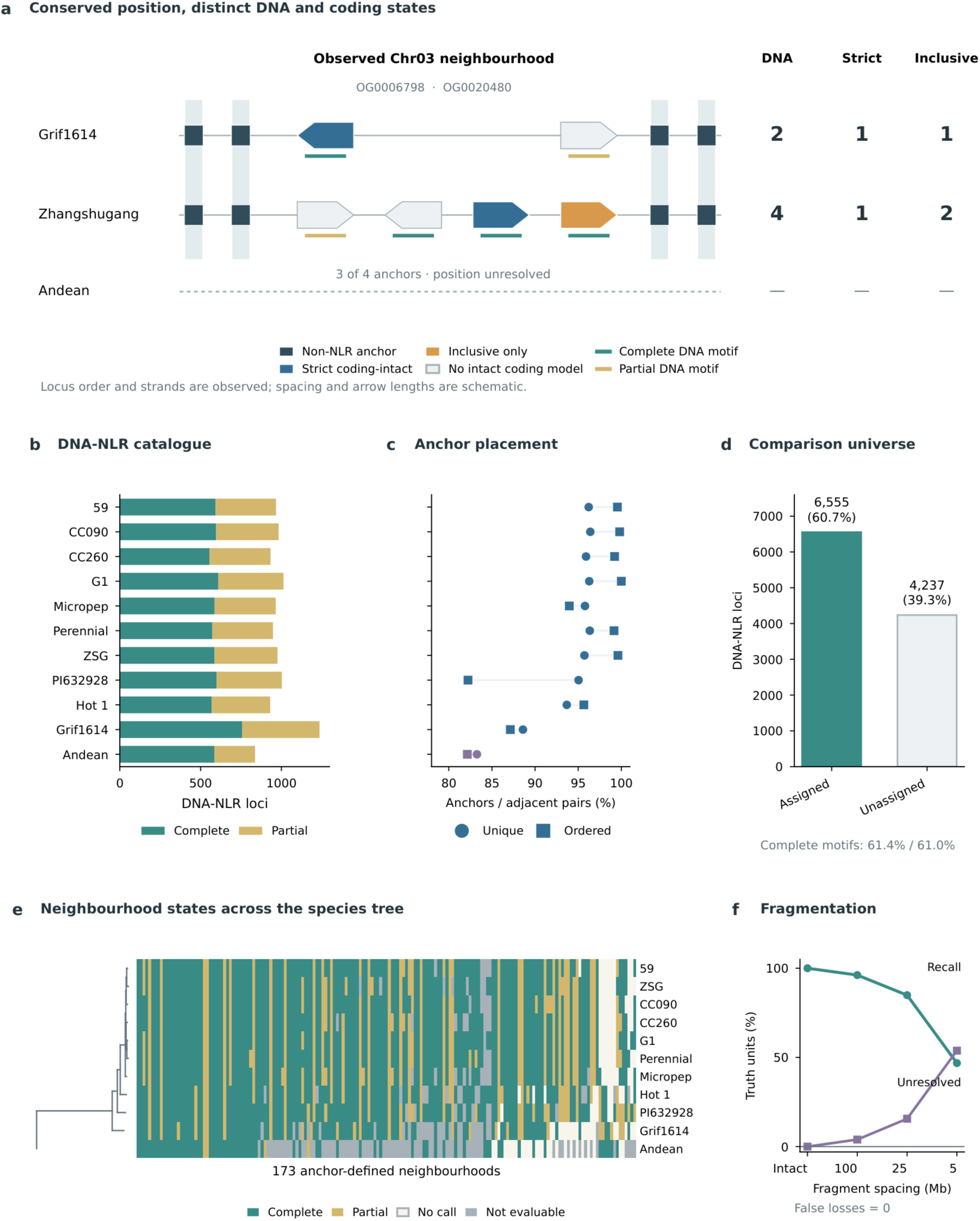
Conserved anchors define an assembly-aware *Capsicum* NLR comparison universe. a, A Chr03 neighbourhood bounded by the OG0006798/OG0020480 inner-anchor pair. Grif1614 has two DNA-NLR calls and one strict/inclusive coding-intact call; Zhangshugang has four DNA calls, one strict call and two inclusive calls. Squares mark conserved non-NLR anchors; arrows show observed NLR order and strand, with schematic lengths and spacing. Arrow fill indicates coding class, and lower marks indicate complete or partial DNA motif architecture. Dashes denote a non-evaluable Andean interval with three of four anchors. b, Complete and partial DNA-NLR architectures across 11 genome assemblies. c, Unique placement of 1,613 non-NLR anchors and adjacent-anchor order consistency. d, Assignment of 6,555 DNA-NLR loci to evaluable neighbourhoods; 4,237 unassigned loci are retained separately. Complete-motif fractions are shown for both groups. e, Species tree and state matrix for 11 genomes and 173 neighbourhoods; grey denotes non-evaluable positions. f, Presence recall and unresolved fractions under simulated fragmentation, with 200 replicates per spacing. The false-loss rate was zero at every spacing. Source data accompany the figure.

We analysed 11 genome assemblies for DNA-level NLR detection and chromosomal comparisons, independently of their annotation availability. A fixed NLR-Annotator workflow identified 10,792 DNA-NLR loci, comprising 6,609 complete and 4,183 partial motif architectures (Fig. 1b; Supplementary Table 1). Per-genome totals ranged from 837 in the Andean *C. rhomboideum* assembly to 1,236 in *C. pubescens* Grif1614. In parallel, 11 representative proteomes contained 382,625 primary proteins and yielded 5,083 protein-NLR candidates. These datasets describe genomic NLR-like sequence and current coding-model support, respectively.

Complete-proteome orthology identified 1,613 stable non-NLR hierarchical orthogroups suitable as positional anchors. Unique genomic placement ranged from 95.7–96.4% among seven *C. annuum* assemblies and from 83.3% to 95.0% in the four more divergent genomes. Adjacent-anchor order consistency was 93.97–100% in *C. annuum* and remained 82.15% in Andean (Fig. 1c; Supplementary Table 2). Three exactly comparable proteomes also showed 90.67–94.72% bidirectional exact-ID concordance with the 2026 pepper pan-NLRome (Supplementary Fig. 1; Supplementary Table 3). This concordance concerns call sets derived from the same public annotations.

Ordered anchors resolved 173 anchor-defined NLR neighbourhoods across the 11 genome assemblies. These units contained 6,555 of the 10,792 DNA-NLR loci (60.7%); the remaining 4,237 loci were retained outside the positional analysis for separate reporting (Fig. 1d). Complete-motif frequency was nearly identical inside and outside the anchor-evaluable set (61.4% and 61.0%), although assigned loci lay farther from assembly edges and N-runs, delimiting the scope of inference to anchor-evaluable NLR neighbourhoods. A maximum-likelihood species tree from 2,069 shared single-copy BUSCO proteins (1,136,749 aligned amino-acid positions) ordered the state matrix. Of the 173 neighbourhoods, 123 were invariantly present among evaluable genomes and 50 were variable; 43 variable units required one transition and seven required repeated transitions. Seventy-five neighbourhoods were evaluable in all 11 genomes, while sample-level evaluability ranged from 99.4% in G1 to 57.2% in Andean. Across the tree, 13 gains and 18 losses were unambiguous and 54 branch placements remained ambiguous across equally parsimonious histories (Fig. 1e; Supplementary Table 4).

Controlled fragmentation quantified how assembly discontinuity affected positional evaluability. Across 200 perturbation replicates, mean presence recall declined from 100% in unbroken anchor shells to 84.9% at 25-Mb and 46.8% at 5-Mb simulated fragment spacing, while 15.6% and 53.8% of units became explicitly unresolved. No fragmentation condition created a false loss (Fig. 1f). Comparisons with annotation synteny, protein orthogroups and position-pure sequence clusters used the same high-confidence anchor-shell reference set, including 19 no-call units (Supplementary Fig. 2).

### DNA-copy expansion recurrently outpaces predicted coding-intact gain

We next compared changes in DNA-copy number with changes in predicted coding-intact copies within anchor-evaluable neighbourhoods. The analysis covered all 173 neighbourhoods and 1,741 positionally evaluable neighbourhood×genome cells, comprising 6,555 distinct DNA-NLR loci (Supplementary Tables 5–8). Standardized leave-one-genome-out recovery, which excluded the target genome annotation, classified 1,428 loci as strict coding-intact and 3,259 under the inclusive definition; 2,304 loci carried an explicit projected coding disruption. Of 1,622 DNA-positive cells, 1,449 (89.3%, neighbourhood-bootstrap 95% CI 85.3–92.9%) contained more DNA than strict coding-intact copies and 1,174 (72.4%, 66.5–78.2%) retained a gap under the inclusive definition. Such gaps occurred in 169 and 156 neighbourhoods, respectively, and were replicated in at least two genomes in 168 and 142 neighbourhoods.

To distinguish a copy deficit from less-than-proportional accumulation, we tested the constant conditional coding-fraction null, which corresponds to an elasticity of one in a log-link count model. Coding-intact copies accumulated more slowly than DNA copies after conditioning on neighbourhood and genome. In the 1,622 DNA-positive cells, Poisson log-link models estimated count elasticities of 0.565 for strict recovery (95% neighbourhood-clustered CI 0.345–0.785; two-sided P = 1.04 × 10⁻⁴ against β = 1) and 0.879 for inclusive recovery (0.779–0.979; P = 0.0181; Figs. 2a and 3a). These estimates correspond to 1.48-fold and 1.84-fold increases in expected coding count per doubling of DNA count. The projection-independent AUGUSTUS/domain route gave β = 0.891 (0.775–1.006; P = 0.0628), a concordant point estimate whose interval includes proportional growth. Complementary coding-fraction models expressed the same association: strict and inclusive odds ratios per DNA-copy doubling were 0.664 (0.544–0.811) and 0.820 (0.694–0.969), respectively (Fig. 2d; Supplementary Tables 9–10).

**Figure 2.**
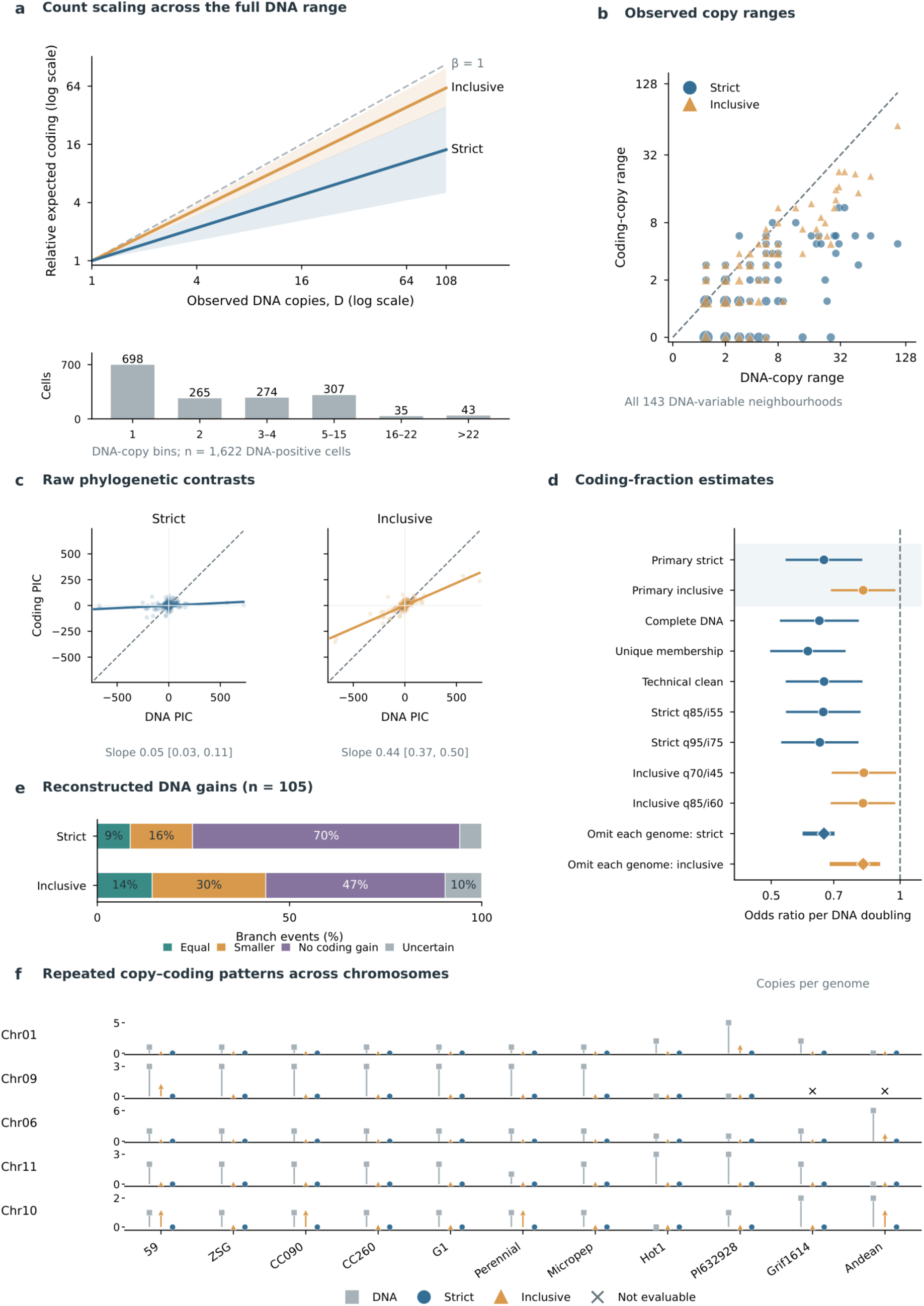
NLR DNA expansion outpaces predicted coding-intact copy accumulation. a, Relative expected coding counts from full-range Poisson models over the observed DNA-copy range (D=1– 108). Curves show Dβ, normalized at D=1; shaded bands transform the two-sided 95% confidence limits for β. Both axes are logarithmic, and the dashed line denotes proportional accumulation (β=1). The histogram shows 698, 265, 274, 307, 35 and 43 DNA-positive neighbourhood×genome cells in bins 1, 2, 3–4, 5–15, 16–22 and >22, respectively (n=1,622). b, DNA-copy range versus strict or inclusive coding-copy range across 143 DNA-variable neighbourhoods. Point area indicates the number of coincident observations, axes use log2(x+1), and the dashed line denotes equal ranges. c, Raw-count regressions through the origin for 573 phylogenetic independent contrasts, with an equal-change reference line. Intervals are 95% neighbourhood-bootstrap confidence intervals for the raw-count slopes. d, Binomial fixed-effect odds ratios for coding-intact fraction per DNA-copy doubling. Circles and horizontal lines show estimates and 95% neighbourhood-clustered confidence intervals; diamonds and thick lines show the primary estimate and the leave-one-genome-out range. e, Joint stepwise-Sankoff classification of 105 unambiguous DNA-gain events into equal coding gain, smaller coding gain, no coding gain and uncertain outcomes. f, DNA, strict and inclusive counts for five prespecified neighbourhoods on different chromosomes, arranged in species-tree order. Each row has its own copy-count scale; crosses denote non-evaluable cells. Directionally unambiguous gains occur in the chromosome 1 and 9 cases.

**Figure 3.**
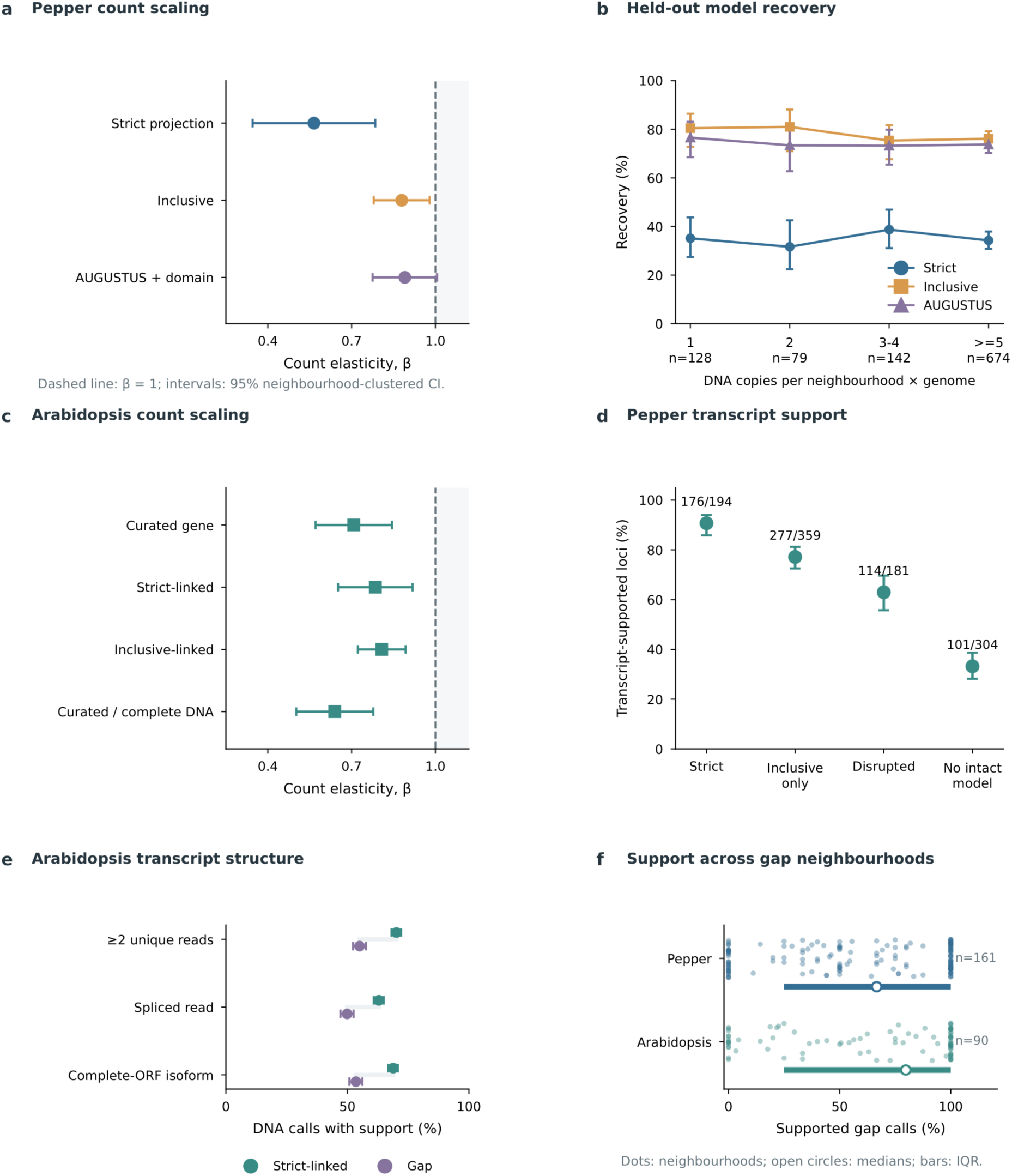
Count scaling, coding-model recovery and transcript support across plant systems. a, Poisson log-link count elasticities for strict, inclusive and projection-independent AUGUSTUS/domain coding definitions in 1,622 DNA-positive pepper neighbourhood×genome cells, including zero coding counts. Points and lines show estimates and two-sided 95% neighbourhood-clustered confidence intervals; the dashed line denotes β=1. b, Recovery of 1,023 held-out complete gene models, with Wilson 95% intervals and bin denominators. c, *Arabidopsis* count elasticities within the published 121-neighbourhood set: curated, strict-linked and inclusive-linked counts against all DNA calls (1,524 cells/112 neighbourhoods), and curated counts against complete-motif DNA calls (1,245 cells/95 neighbourhoods). d, Transcript-support proportions in matched G1 and Andean RNA-seq, with Wilson 95% intervals; labels report supported/evaluated loci. e, *Arabidopsis* support from at least two MAPQ ≥20 locus-unique reads, a locus-unique spliced read or a locus-unique complete-ORF transcript model. Denominators are 2,105 strict-linked and 1,316 gap DNA calls; intervals are Wilson 95% intervals. f, Transcript-supported fractions among gap calls in eligible neighbourhoods. Each dot represents a neighbourhood; open circles mark medians and thick lines show interquartile ranges. Vertical jitter separates overlapping observations.

Coarse copy-number bins did not show a monotonic decline in held-out recovery, whereas finer bins resolved lower strict recovery in the high-copy tail. Among 1,023 complete published models excluded from target-state calling, strict recovery was 35.2%, 31.6%, 38.7% and 34.3% in cells with 1, 2, 3–4 and ≥5 DNA copies; projection-independent recovery was 76.6%, 73.4%, 73.2% and 73.7% (Fig. 3b). DNA copy number did not predict strict or projection-independent recovery after genome or locus-length adjustment. Inclusive recovery showed a modest neighbourhood-adjusted decline. Binomial null simulations preserving neighbourhood and genome effects, including the empirical copy-bin recovery profile, did not reproduce the observed coding-fraction odds ratio. In finer bins, strict recovery was 144/357 (40.3%) at 5–15 DNA copies, 22/69 (31.9%) at 16–22 and 65/248 (26.2%) above 22. The >22-versus-5–15 contrast adjusted for genome and locus length gave an odds ratio of 0.566 (95% neighbourhood-clustered CI 0.290–1.102); the interval permits a substantial recovery decline (Supplementary Table 11).

The strict count relationship persisted across the principal sequence and sampling controls. Its elasticity was 0.571 (0.282–0.860) among the 75 neighbourhoods evaluable in all 11 genomes, 0.532 (0.327–0.736) in exact four-anchor cells and 0.654 (0.463–0.845) after excluding DH06. All seven species-balanced fits and all 11 leave-one-genome-out fits had 95% intervals below one (Fig. 3a; Supplementary Fig. 3; Supplementary Table 10). The strength of the relationship depended on the sampled copy-number range: restricting cells to at most 15 or 22 DNA copies moved strict elasticities to 0.845 (0.607–1.084) and 0.887 (0.683–1.091). Thus, highly expanded arrays contribute substantially to the full-range effect. Inclusive and AUGUSTUS sensitivities also retained point estimates below one, with some intervals spanning one. Omission of each pepper neighbourhood retained 95% intervals below one in all 173 strict and all 173 inclusive fits; the complete influence ranges and high-copy recovery diagnostics are reported in Supplementary Tables 11–12.

The count association persisted when the analysis was restricted to complete DNA evidence. Among 4,023 complete-motif pepper loci, strict and inclusive count elasticities were 0.591 (95% CI 0.377– 0.805) and 0.876 (0.808–0.944). Requiring unique neighbourhood membership and an N fraction ≤0.01 retained 3,890 loci and gave β = 0.536 (0.314–0.759) and 0.863 (0.801–0.925), respectively (Supplementary Tables 10 and 13–15). The corresponding raw-range and branch summaries also retained widespread DNA–coding disparity (Supplementary Fig. 4).

Observed ranges and reconstructed histories described the resulting copy disparity. Among 143 DNA-variable neighbourhoods, coding range was smaller than DNA range in 115 strict and 96 inclusive comparisons, whereas five and ten showed the reverse (Fig. 2b). Across 573 phylogenetic independent contrasts, raw through-origin coding-on-DNA slopes were 0.049 (95% neighbourhood-bootstrap CI 0.025–0.106) and 0.437 (0.365–0.498); the partitioned BUSCO tree gave similar estimates (Fig. 2c). Joint parsimony reconstruction identified 105 unambiguous DNA-gain events. Ninety strict and 80 inclusive events had smaller or absent coding gains, and 77 met this definition under both coding criteria (Fig. 2e). These summaries describe numerical copy changes; the count and fraction models above test the constant-fraction hypothesis.

A DH06-excluded screen identified five replicated loci on chromosomes 1, 6, 9, 10 and 11 (Fig. 2f; Supplementary Tables 16–18). The chromosome 1 case expanded to five DNA copies in PI 632928 but retained one inclusive predicted coding-intact copy, against a widespread one-copy background. The chromosome 9 case contained three DNA copies throughout the *C. annuum* clade but zero or one inclusive predicted coding-intact copy, while Hot1 and PI 632928 carried no DNA copy. A distinct chromosome 6 neighbourhood reached six DNA copies in Andean, of which one was inclusive predicted coding-intact and four were explicitly disrupted. The chromosome 11 and 10 cases independently combined two-to-three DNA copies with zero-to-one inclusive predicted coding-intact copies across different lineages. Only the chromosome 1 and 9 gains were directionally unambiguous across all equally parsimonious branch histories, so the other three are used as replicated observed patterns rather than polarized gain claims. Repeating the complete recovery workflow with 25-kb rather than 12-kb flanks gave exact strict and inclusive calls at all 94 selected-case loci and retained all five frozen-screen cases (Supplementary Fig. 4; Supplementary Table 19), excluding extraction-window width as an explanation. The copy–coding patterns and available structural support are summarized in Supplementary Table 20.

### Sublinear count scaling recurs in *Arabidopsis* and coding states differ in transcript support

Sublinear coding-copy accumulation recurred in the published 121 graph-defined NLR neighbourhoods from 17 *A. thaliana* accessions (Teasdale et al. 2025). Curated-gene counts had an elasticity of 0.708 (95% neighbourhood-clustered CI 0.571–0.844; two-sided P = 2.80 × 10⁻⁵), while strict-linked and inclusive-linked counts gave 0.785 (0.651–0.918) and 0.808 (0.723–0.893), respectively (Fig. 3c). The primary model included 1,524 DNA-positive accession×neighbourhood cells spanning 112 neighbourhoods; zero-DNA cells remained in the released matrix and were outside the log-DNA estimand. Curated annotations and DNA calls were not perfectly nested: 96 evaluable cells contained curated genes without DNA calls, and 107 DNA-positive cells contained more curated genes than DNA calls (Supplementary Table 21). The linked-count analysis directly addresses this counting mismatch. Curated-gene elasticities remained below one after excluding the five largest neighbourhoods (0.603, 0.418–0.789), in every leave-one-neighbourhood-out fit (range 0.671–0.751) and in every leave-one-accession-out fit (0.679–0.727; Supplementary Table 10). Neighbourhood-level range attenuation was not reproduced, and some curated-status sensitivities included proportional growth; the shared finding is the conditional count association.

Matched transcript data allowed us to compare RNA support across genomic coding states. Across G1 and Andean, strict RNA-seq support was detected for 176 of 194 predicted strict-intact loci (90.7%), 277 of 359 inclusive-only loci (77.2%), 114 of 181 explicitly disrupted loci (63.0%) and 101 of 304 loci without an intact coding model (33.2%; Fig. 3d). After adjustment for locus length, neighbourhood DNA copy number and matched genome, strict-intact loci retained 10.7-fold higher odds of transcript support than loci without an intact model (95% CI 6.29–18.1); inclusive-only and disrupted loci retained 4.09-fold (2.64–6.32) and 2.43-fold (1.61–3.67) higher odds. The gradient also remained in locus-by-tissue models. Accession-matched *Arabidopsis* Iso-Seq was then resolved beyond interval overlap into locus-unique reads, splice-junction-bearing reads, collapsed transcript structures and complete predicted ORFs (Fig. 3e). Locus-unique spliced reads supported 1,325 of 2,105 coding-linked calls (62.9%) and 656 of 1,316 copy–coding-gap calls (49.8%); locus-unique complete-ORF transcript models supported 1,449 (68.8%) and 704 (53.5%), respectively. After adjustment for locus length, neighbourhood DNA-copy number and accession, no association of complete-ORF transcript support with the coding-link boundary was detected (odds ratio 1.19, 95% CI 0.78–1.82), nor was an association detected for locus-unique spliced reads (1.02, 0.67–1.56). These intervals do not establish equivalence between coding classes. Across all eligible neighbourhoods rather than selected cases alone, gap-copy support ranged from none detected to support for every gap call (Fig. 3f). Expanded NLR-like DNA pools therefore combine predicted intact copies, non-intact copies with locus-level transcript structures and copies without detected transcripts in the sampled libraries.

### Neighbourhood turnover occurs in a repeat-rich structural background

The G1–Andean comparison contained 78 neighbourhoods evaluable in both genomes and 30 state-contrast units with paired interval and repeat measurements. Outgroup polarization with tomato, potato, *C. chacoense* and *C. galapagoense* resolved both Andean-lineage losses and non-Andean *Capsicum* gains, showing that the same extant contrast arose in multiple directions (Supplementary Fig. 5; Supplementary Tables 22–23). Repeat differences tracked some interval-length contrasts, but matched four-anchor NLR-free controls showed no NLR-neighbourhood-specific excess: the median neighbourhood-minus-control repeat expansion was −0.50 percentage points. A common-library Dfam rerun preserved the broad direction of interspersed-repeat and Gypsy contrasts.

### DH06 preserves an anchor shell while internal NLR families are remodelled

DH06 provided the most resolved structural example. Local whole-genome alignment and a reciprocal-position graph connected 11 G1 families, F392–F402, across five representative genomes within a conserved non-NLR anchor shell (Fig. 4a). Seven *C. annuum* assemblies retained one copy of all 11 families in collinear order. PI 632928 and Hot 1 showed family-specific replacement, loss or duplication. Grif1614 expanded selected families within a 17.35-Mb broad interval, whereas the Andean interval contracted to 0.55 Mb and contained no DNA-NLR call.

**Figure 4.**
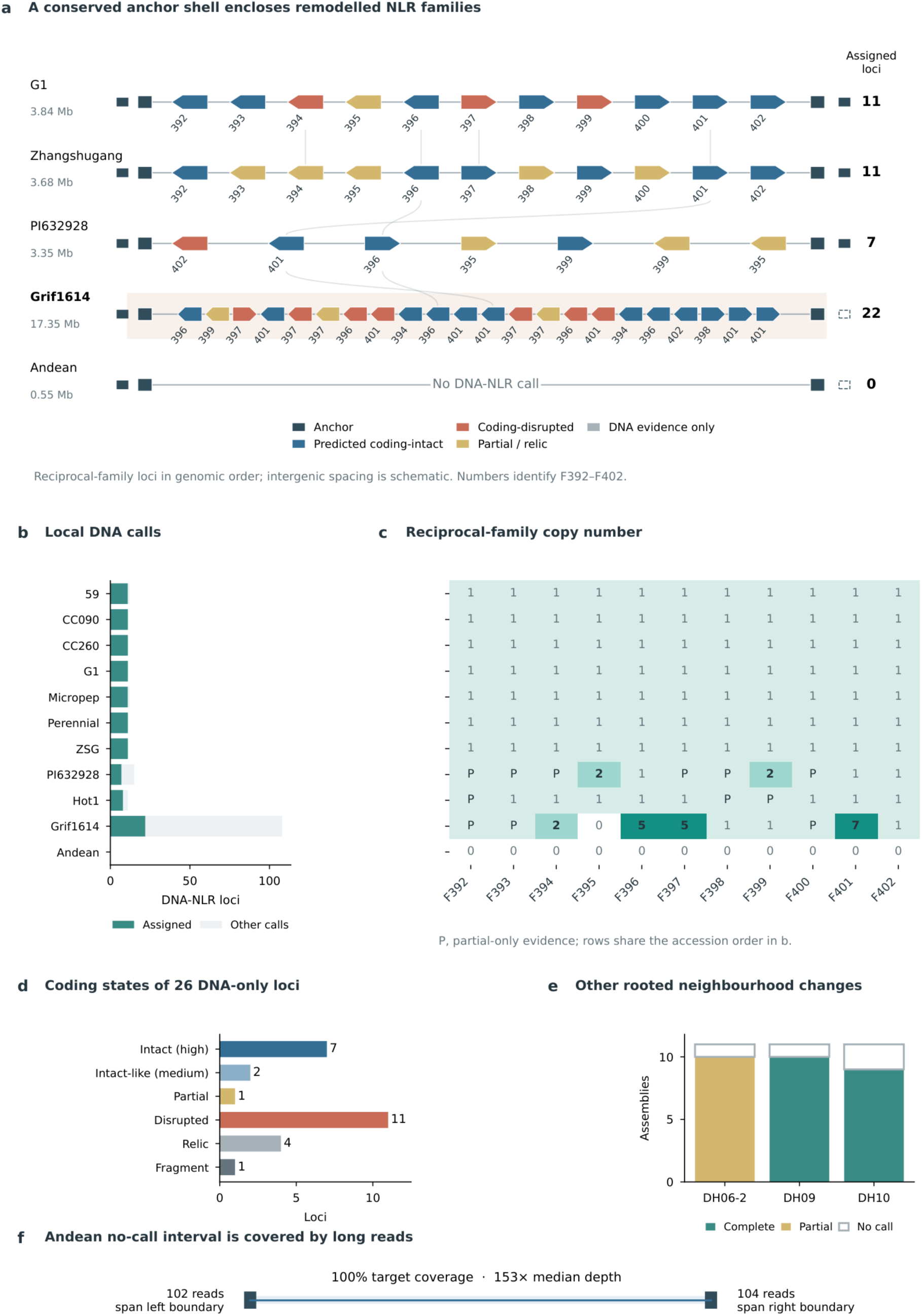
DH06 retains a conserved anchor shell while its NLR families are repeatedly remodelled. a, Reciprocal-family loci in observed genomic order within conserved non-NLR anchors. Arrow direction indicates strand and fill denotes predicted coding state; labels 392–402 identify families F392–F402. Spacing and arrow lengths are schematic, with the full anchor-interval length printed for each genome. Curves connect family-level median positions for F394, F396, F397 and F401. Open outer-anchor marks indicate the absence of a unique primary projection. b, All local DNA-NLR calls and the subset assigned to F392–F402. c, Family-copy matrix across 11 genomes; P denotes partial-only evidence. d, Coding-state classification of 26 DNA-only reciprocal-family loci. e, Three rooted neighbourhood changes outside DH06. f, Schematic of the Andean validation interval showing observed coverage, median depth and reads spanning each inner-anchor boundary. On Zhangshugang coordinates, the F392–F402 core lies 1.32 Mb distal to the Chr06 introgression interval reported by Liu et al. (2023), whereas the broad anchor shell overlaps its distal edge (Supplementary Table 24).

The distinction between density and positional homology was especially important in Grif1614. Its broad DH06 interval contained 108 DNA-NLR loci, but only 22 were assigned to the 11 reciprocal-position families; 86 remained unassigned or ambiguous (Fig. 4b,c). F394, F396, F397 and F401 were duplicated or expanded among assigned loci. The Andean interval retained the anchor shell but none of the 11 families. Competitive whole-genome mapping of Andean PacBio reads covered 100% of the 751-kb validation target at a median depth of 153×, with 102 and 104 clean reads spanning the left and right inner-anchor edges (Fig. 4f). Continuous read support across the contracted interval supports an assembled, callable local contraction rather than a gap.

Published annotations supported only a subset of assigned DH06 loci. We therefore evaluated 26 DNA-only reciprocal-family loci using family-matched protein projection, de novo gene prediction, NB-ARC profile search and NLRtracker. Seven were classified as high-confidence predicted coding-intact NLRs, two as medium-confidence intact-like, one as partial, 11 as coding-disrupted, four as conserved relics and one as a fragment (Fig. 4d). The rescued models show that annotation-only comparisons miss plausible intact coding sequences and obscure differences among DNA copies.

### A near-megabase duplication contains a copy-specific coding-state mosaic

The Grif1614 expansion contained two same-orientation blocks of 0.931 and 0.946 Mb on chromosome 6. Eighteen collinear segments comprised 866,550 matched bases at 99.5804% aggregate identity, identifying a recent direct segmental duplication (Fig. 5a). Five copy-specific structural events were grouped into four composition units for plotting: a 35.833-kb copy-2 gain, two adjacent copy-2 components of 8.423 and 6.645 kb, a 2.299-kb copy-1 compensating sequence and a 10.344-kb distal copy-1 remodelling event (Fig. 5a,d). Their repeat compositions differed sharply; no single repeat class, including Ale/Gypsy, explained all boundaries. Each of 19 tested structural edges was spanned by 6– 24 clean MAPQ ≥20 HiFi reads; the minimum within each of the six event groups ranged from 6 to 13 (Fig. 5e).

**Figure 5.**
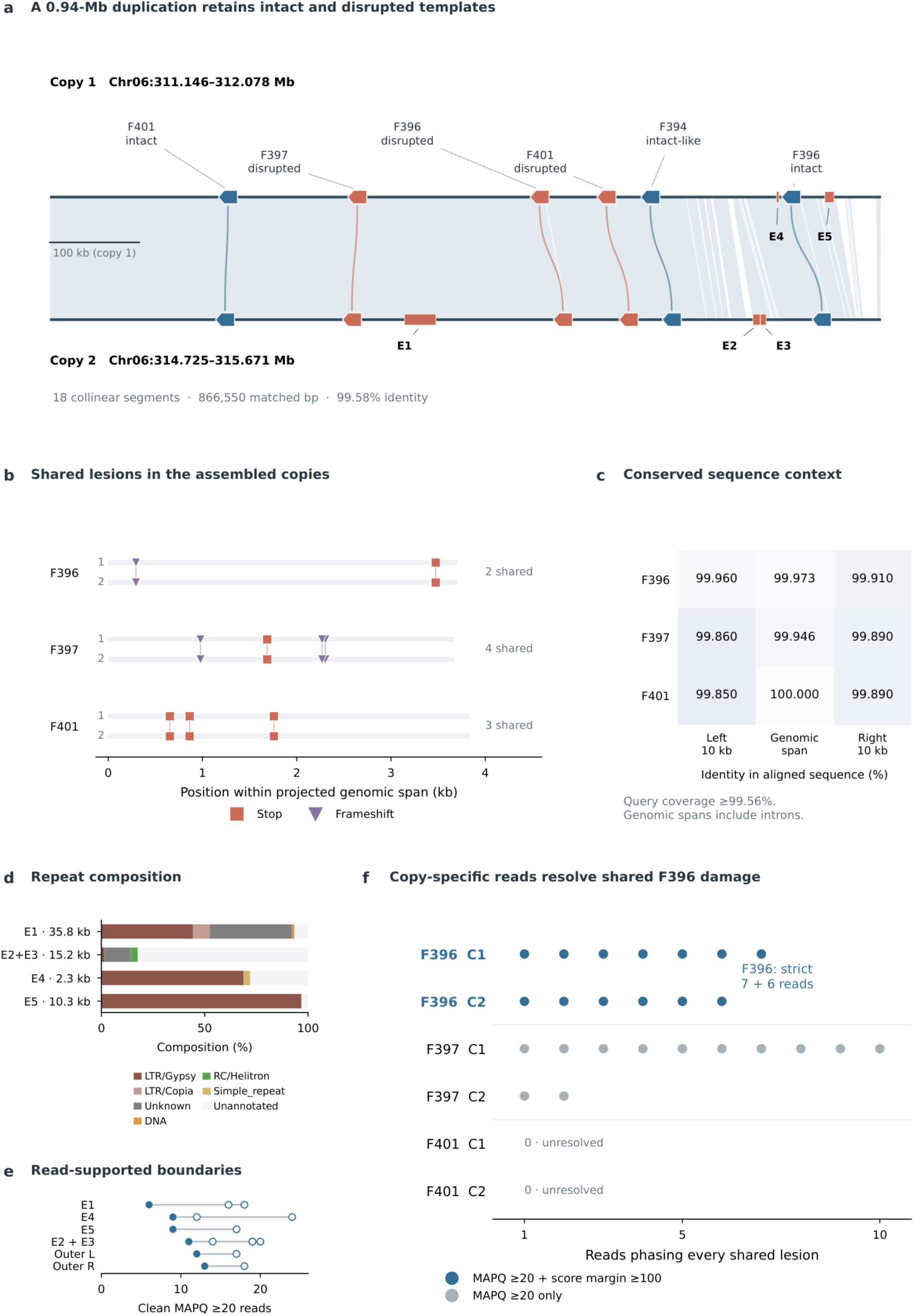
A recent block duplication contains a copy-specific mosaic of predicted coding states. a, Same-orientation alignment of two Grif1614 blocks, showing genomic coordinates, six family pairs and structural events E1–E5. Arrow direction denotes strand; blue and terracotta indicate predicted intact/intact-like and disrupted states. Pale polygons show 18 alignment segments, curves connect family pairs, and rectangles mark copy-specific sequence differences. E2 and E3 form one adjacent composition unit. b, Shared stop and frameshift positions in assembled copies 1 and 2 for F396, F397 and F401. c, Sequence identity across projected genomic spans and 10-kb flanks. Values are percentages; genomic spans include introns. Query coverage is 100% except for the F396 right flank (99.56%). d, Repeat composition of four units representing the five structural events. e, Clean MAPQ ≥20 read support across 19 tested structural boundaries. Open dots show individual edges, lines span the observed range, and filled dots mark group minima. Counts ranged from 6 to 24; Outer L/R denote the duplication boundaries. f, Copy-specific MAPQ ≥20 reads phasing all assayed shared lesions at each locus. Blue dots additionally satisfy a ≥100-point best-to-competing alignment-score margin. F396 is supported under the strict criterion, F397 under the broader criterion, whereas F401 remains unresolved at the physical-copy level. Source data are provided in Supplementary Tables 25–26.

Six reciprocal-family pairs connected the two duplicated blocks, including intact or intact-like F394, F396 and F401 pairs and disrupted F396, F397 and F401 pairs. Five had ≥99.5% query coverage in their 100-kb windows; the intact F396 pair occupied a more remodelled window (86.3% coverage; Supplementary Table 26). The three disrupted pairs shared two, four and three lesion signatures, respectively, with no private lesion among the assayed events (Fig. 5b). Base-level alignment of their projected genomic spans revealed one nucleotide difference across 3,703 bp in F396, two across 3,670 bp in F397 and none across 3,834 bp in F401. Their left and right 10-kb flanks also retained 99.85– 99.96% identity over aligned sequence, with 99.56–100% query coverage (Fig. 5c; Supplementary Table 26). The shared coding lesions therefore occur within broadly conserved duplicated sequence.

The raw reads differed in how well they assigned damaged templates to physical copies. Seven copy-1 and six copy-2 F396 reads each phased both lesions at MAPQ ≥20 and a ≥100-point best-to-competing alignment-score margin. Ten copy-1 and two copy-2 F397 reads phased all four lesions at MAPQ ≥20, but none met the strict score margin. F401 had exact event-level support without reads assigning all three lesions to a physical copy (Fig. 5f; Supplementary Table 25). F396 thus establishes that both physical blocks contain the same damaged coding template; F397 supports this interpretation at moderate confidence and F401 remains unresolved. This is consistent with amplification of pre-existing damaged templates, with later inter-copy gene conversion as an alternative to that chronology.

A DH06-derived junction module also segregated among 38 *C. pubescens* accessions (Supplementary Results, “Population context of the DH06-derived cross-copy module”; Supplementary Figs. 6–7; Supplementary Table 27).

## Discussion

NLR expansion changes both array size and coding composition. Across the surveyed *Capsicum* neighbourhoods, predicted coding-intact counts rose less than proportionally with DNA counts after accounting for neighbourhood and genome effects; the conditional relationship recurred in the independently defined *Arabidopsis* panel. This result complements the established view of NLR repertoires as products of duplication, diversification and degradation (Michelmore and Meyers 1998; Teasdale et al. 2025). The quantitative distinction is that larger DNA-defined arrays do not simply contain proportionally more recoverable intact coding copies. At the level of physical copies, the DH06 duplication demonstrates that two physical blocks retain the same disrupted F396 state alongside intact family pairs. The comparative and local results thus describe different aspects of what expanded NLR-associated DNA retains.

The proportionality test asks more than whether genes and pseudogenes coexist. Earlier studies established uneven expansion within NLR clusters and substantial pseudogene content in variable repertoires (Lee and Chae 2020; Winters et al. 2025). Neither observation alone requires coding fractions to decline with DNA-copy number. Our count models test this relationship, while complementary fraction models describe the associated change in composition. The inclusive pepper estimate corresponds to a 1.84-fold increase in expected coding count for a doubling of DNA count: accumulation is attenuated, not absent. The independently curated *Arabidopsis* counts and their DNA-linked counterparts recover the same direction without importing the pepper neighbourhood definition. Cross-system agreement therefore concerns conditional count scaling.

Several evolutionary histories could generate the observed composition, including disruption after duplication, amplification of already disrupted sequence, and differences among copies in persistence or loss. Sequence exchange is another possibility: experiments have documented the roles of unequal recombination and gene conversion in resistance-gene cluster evolution, with incidence varying among members of the same locus (Chin et al. 2001; Kuang et al. 2004). Fitness costs demonstrated for *Arabidopsis* RPM1 make selection on receptor maintenance a plausible additional influence (Tian et al. 2003). Distinguishing that explanation from neutral turnover and heterogeneous constraint requires evidence beyond the present count comparisons.

DH06 narrows the inference from an array-level count difference to the state of identifiable physical copies. The megabase-scale organization of this locus has precedents in resistance-gene clusters (Meyers et al. 1998), whereas block alignment and copy-specific lesion phasing resolve which coding states occupy the duplicated sequence. Seven copy-1 and six copy-2 F396 HiFi reads each support both shared lesions under the strict assignment criterion, within broadly conserved duplicated sequence. Both physical blocks therefore carry the same disrupted coding state. Duplication of a previously damaged template is a parsimonious history, whereas subsequent inter-copy gene conversion remains compatible with the data. Copying a mixed block can preserve its coding fraction. DH06 therefore demonstrates the physical coexistence of intact and disrupted templates after expansion, but does not by itself explain the sublinear relationship across neighbourhoods. The five additional chromosomal cases extend the comparative recurrence of copy–coding patterns.

Transposon proliferation has shaped pepper genome architecture, and retroduplication has contributed to its resistance-gene repertoire (Kim et al. 2017; Zhang et al. 2025). Our matched NLR-free intervals showed no neighbourhood-specific excess of repeat expansion, while the DH06 structural events had distinct repeat compositions. Repeat-rich sequence therefore provides a heterogeneous structural background for the observed remodelling.

Transcript support further characterizes the copies within expanded arrays. In matched pepper libraries, support was most frequent for strict coding-intact predictions but was also detected at explicitly disrupted loci. *Arabidopsis* Iso-Seq recovered spliced structures and complete transcript ORFs on both sides of the coding-link boundary; these data also contributed to the original annotations. Established NLR regulation illustrates why genomic and transcript states can differ: RPS4 resistance depends on both regular and alternatively spliced transcripts, and FPA-mediated premature transcription termination regulates numerous *Arabidopsis* NLRs (Zhang and Gassmann 2003; Parker et al. 2021). These precedents provide biological context for the observed transcript diversity; the mechanisms at individual pepper loci remain to be established.

Noncanonical architecture also warrants biological attention. The atypical TIR-NBS2 protein, which lacks the LRR domain, is required for activated defence in the *Arabidopsis* exo70B1 mutant (Zhao et al. 2015). NLR output can depend on sensor–helper relationships, as demonstrated for the NRC network (Wu et al. 2017). Coding integrity is consequently one component of candidate evaluation alongside domain architecture, transcript support and genomic context. Our classification retains explicit lesions, incomplete architecture and insufficient recovery evidence as distinct states.

Measurement and sampling limit the strength and scope of the evolutionary inference. Only 60.7% of pepper DNA-NLR loci entered anchor-evaluable neighbourhoods, so the relationship should not be extrapolated to unassigned loci. Omission of every individual neighbourhood retained confidence intervals below one for strict and inclusive pepper counts, arguing against a single-neighbourhood explanation. This does not remove dependence on the high-copy tail as a group. Strict held-out recovery fell from 40.3% at 5–15 DNA copies to 26.2% above 22, and the adjusted contrast remains compatible with a substantial recovery decline. Moreover, the projection-independent estimate and upper-tail-restricted fits include proportionality in their intervals. Thus, the available checks support a recurrent relationship in the reported counts but do not separate biological attenuation completely from heterogeneous coding recovery. The two-system comparison is evidence of recurrence, not a universal plant scaling law or a calibrated estimate of functional receptor dosage.

These distinctions matter when NLR repertoires are used to identify resistance candidates. Resistance gene enrichment sequencing (RenSeq) supports NLR discovery and genetic mapping, while association genetics combined with RenSeq connects sequence variation to resistance phenotypes (Jupe et al. 2013; Arora et al. 2019). Our results motivate coding-state-aware interpretation of these resources and of NLR candidates within genome-wide association or quantitative trait locus intervals. Alongside total DNA-copy number, candidate summaries should retain predicted coding-intact counts, explicit coding lesions, unresolved models and locus-specific transcript support. Candidates with different evidence profiles can then be prioritized separately, rather than treating every additional DNA call as an interchangeable receptor. This is an evidence-stratification proposal, not a validated numerical weighting scheme: the present study estimates neither copy-specific functional weights nor improvements in resistance prediction.

By quantifying coding composition within homologous arrays, this study complements gene-centred pan-NLRomes, positional comparisons and analyses of sequence diversity (Prigozhin and Krasileva 2021; Dong et al. 2026; Guo et al. 2025). It pairs a direct proportionality test across two plant systems with a copy-resolved example of shared coding damage. Copy-resolved transcript and protein measurements can connect these genomic states to receptor products, while genetic evidence can establish their contributions to immunity.

## Methods

### Genome, annotation and read resources

The genome-assembly panel comprised seven *C. annuum* accessions (59, CC090, CC260, G1-36576, Micropep Red, Perennial and Zhangshugang), *C. baccatum* PI 632928, *C. chinense* Hot 1, *C. pubescens* Grif1614 and *C. rhomboideum* Andean. Assembly accessions, source databases, checksums and sequence-role classifications are provided in Supplementary Table 1. The annotation/proteome panel contained 11 representative proteomes; a primary transcript was selected from annotation tags where present, then by CDS length, protein length and stable transcript identifier. UCD-10X-F1 was retained only as a sensitivity resource and excluded from independent-lineage counts. Public resource context followed the original genome studies (Liu et al. 2023; Chen et al. 2024; Zhang et al. 2025).

Tomato, potato, *C. chacoense* CGN21477 (GCA_053540505.1) and *C. galapagoense* CGN22208 (GCA_053540595.1) supplied outgroup intervals. Grif1614 PacBio reads were obtained from SRR23352910 and Andean reads from SRR24319436. Population analyses used 38 public *C. pubescens* paired-end datasets from PRJNA801499.

### DNA- and protein-level NLR evidence

NLR-Annotator v2.1b (commit 5d7f1ecfae8927f34baa1126998ccc9869995784) was run on each of the 11 assemblies with fixed distances of 500 bp within motif combinations, 2,500 bp for elongation and 10,000 bp between motif combinations (Steuernagel et al. 2020). Calls were normalized to BED coordinates and retained as complete or partial motif architectures; stop-containing loci and non-primary sequence roles were labelled rather than removed. Protein candidates were classified with NLRtracker v1.0.3 and InterProScan 5.53-87.0 (Kourelis et al. 2021). Isolated NB-ARC-only proteins were retained in an exclusion table. DNA loci and gene models were reconciled by genomic overlap, using 50% of the shorter feature as the primary threshold. Evidence states were complete-model supported, partial-model supported, DNA-only, protein-only or not evaluable. A predefined 491-case evidence audit included 100 positive controls, 340 stratified DNA loci, 16 conflict cases and 38 protein-only cases, with three overlaps between strata; every coordinate, identity, threshold, strand and relation-accounting claim was independently recomputed from the underlying tables before closure.

### Stable anchors and neighbourhood construction

OrthoFinder v3.1.5 with DIAMOND v2.2.4 was applied to all representative proteomes in the annotation panel (Emms and Kelly 2019; Buchfink et al. 2021). Anchor HOGs were required in at least eight proteomes, single copy in at least seven and no more than two copies in any proteome. NLR-associated groups and unstable length outliers were excluded. The 1,613 retained HOGs were projected into all 11 genome assemblies with miniprot v0.18-r281 (Li 2023). A primary placement required query coverage ≥0.50 and a best-to-second score ratio ≥1.05. Unique placement, sequence role, orientation and adjacent-anchor order were recorded independently.

Each DNA-NLR observation received an ordered signature from two inner and two outer anchors where available. Exact four-anchor signatures formed the strict class. A unique compatible three-anchor signature was accepted when order and orientation were continuous; other non-exact high-confidence assignments required local whole-genome alignment. Multiple loci from the same genome could be assigned to the same neighbourhood but were retained as distinct copy observations. Each neighbourhood was back-projected into all 11 assemblies. A completely scanned, uniquely delimited, continuously assembled interval without a DNA-NLR call was labelled NO_NLR_CALL; broken, ambiguous or multi-position intervals were NOT_EVALUABLE.

Released tables retain their original stable neighbourhood keys, including PNOU_DH06_N0OG0010401_N0OG0019490_001; these identifiers link observations across files. The manuscript uses the term “neighbourhood”.

### Target-annotation-independent coding-state recovery across all neighbourhoods

For every DNA-NLR locus assigned to a neighbourhood, we extracted the locus plus 12 kb on each side and recovered coding models without consulting the target genome’s published annotation. Protein queries were drawn from high-quality NLR candidates in the other ten genomes within the same neighbourhood; all same-genome queries were excluded. Amino-acid-aware projection used miniprot, and an independent AUGUSTUS tomato-parameter prediction was evaluated by HMMER against PF00931 and by NLRtracker. A projected or predicted feature had to overlap the DNA-NLR locus by at least 50% of the shorter feature. The strict call required an other-genome high-quality projection with query coverage ≥0.90, amino-acid identity ≥0.65, no projected frameshift, in-frame stop or internal translation stop, and an NLRtracker NLR classification. The inclusive call comprised the strict set plus either an intact other-genome projection with coverage ≥0.75 and identity ≥0.50, or an AUGUSTUS protein ≥500 aa with PF00931 model coverage ≥0.75; both routes also required NLRtracker NLR support. A projection with coverage ≥0.75, identity ≥0.50 and at least one explicit coding disruption was recorded separately as coding-disrupted. Loci lacking sufficient evidence remained partial, divergent or unresolved rather than being called coding-intact.

Locus calls were summed within each of the 173 neighbourhood×11-genome cells. Positional non-evaluability remained missing, whereas an evaluable neighbourhood with no DNA-NLR locus retained a biological zero. Pre-specified sensitivities varied strict projection coverage/identity to 0.85/0.55 and 0.95/0.75, inclusive projection thresholds to 0.70/0.45 and 0.85/0.60, and AUGUSTUS length/PF00931 coverage to 400 aa/0.65 and 600 aa/0.85. Further analyses retained only complete-motif DNA loci, uniquely assigned loci, technically clean windows at three N-content thresholds, or excluded Andean. The projection-independent route counted an AUGUSTUS model only when it was complete before domain filtering and received both PF00931 and NLRtracker NLR support.

Recovery sensitivity was evaluated against 1,023 COMPLETE_MODEL_SUPPORTED loci with unique neighbourhood membership. These target annotations defined the held-out positive set but were not available to the recovery routes. Sensitivity was summarized in neighbourhood×genome DNA-copy bins of 1, 2, 3–4 and ≥5 with Wilson intervals. Cluster-robust logistic models tested copy number while adjusting for genome and, separately, neighbourhood or locus length. The earlier 26-locus DH06 rescue remained an independent local calibration of strict and inclusive classification boundaries.

### Positional evaluability under fragmentation

Controlled fragmentation tested whether loss of anchor continuity was classified as non-evaluable rather than biological absence. Full perturbation parameters and scoring rules are provided in Supplementary Methods, under “Detailed supporting analyses”.

### Species tree and phylogeny-aware turnover

Single-copy BUSCO v6.1.0 proteins shared across all 11 genome assemblies were obtained with the eudicots_odb10 lineage and aligned individually with MAFFT v7.526 (Manni et al. 2021; Katoh and Standley 2013). All 2,069 shared loci were retained; alignment columns containing a non-gap, non-ambiguous amino acid in at least eight genomes were concatenated into a 1,136,749-aa supermatrix. A maximum-likelihood tree was inferred in IQ-TREE v2.4.0 under LG+F+G4 with 1,000 ultrafast bootstrap and 1,000 SH-aLRT replicates, then rooted with Andean (Minh et al. 2020). The unpartitioned concatenated model supplied the primary species-tree ordering; the same locus set was also analysed with 2,069 partitions as a topology sensitivity analysis.

For each of 173 neighbourhoods, complete and partial DNA-NLR states were coded present, an evaluable no-call was coded absent and NOT_EVALUABLE was treated as missing. Equal-cost binary Sankoff reconstruction enumerated optimal parent–child state pairs. A branch was called an unambiguous gain or loss only when every optimal reconstruction required the same transition; otherwise placement was ambiguous. Units with fewer than four evaluable genomes were labelled insufficient. Remaining units were invariant present, invariant absent, single-transition or repeated-transition.

### Copy–coding relationships within evaluable neighbourhoods and replicated-case analysis

The primary count model was a Poisson generalized linear model with a log link, log E(C_ng) = alpha_n + gamma_g + beta log(D_ng), where C and D denote coding and DNA counts and n and g index neighbourhood and accession. It was fitted to positionally evaluable cells with D>0 and retained C=0. Neighbourhood and accession were fixed effects; uncertainty was estimated using the neighbourhood-clustered sandwich covariance. At β=1 the expected coding fraction is constant conditional on these effects; β<1 denotes sublinear coding-copy accumulation. We report two-sided 95% Wald intervals and two-sided tests against β=1, with directional probabilities supplied separately in source data. Expected coding-count and coding-fraction changes per DNA doubling are 2β and 2(β−1). Models were repeated in fully evaluable neighbourhoods, exact four-anchor cells, cells with D≤15 or D≤22, species-balanced panels, leave-one-accession-out panels, the DH06-excluded set, and the stated sequence and coding-definition sensitivities. The earlier log1p-on-log1p coefficients remain descriptive transformed-count associations in source data; a slope of one on that scale is not a general constant-fraction null.

Second, among DNA-positive cells, a binomial model used the number of predicted coding-intact and non-intact DNA copies as the grouped response, log2 DNA copy number as the focal predictor, and neighbourhood and genome fixed effects, with neighbourhood-clustered sandwich intervals. The exponentiated coefficient is the odds ratio for the coding-intact fraction per doubling of DNA dosage. A null simulation fitted coding probability without a copy-number term while retaining neighbourhood and genome effects, generated 1,000 binomial datasets and refitted the full model. A second 1,000-replicate simulation additionally imposed the empirical held-out recovery rates from the four DNA-copy bins. Calling-threshold and leave-one-genome-out sensitivities retained the same model structure.

DNA and coding counts were also summarized by maximum-minus-minimum ranges and phylogenetic independent contrasts after pruning each neighbourhood to evaluable tips. Through-origin raw-count slopes and 10,000 neighbourhood-bootstrap intervals describe the contrast relationship. Equal raw ranges, PIC slope one and equal branch increments represent equal numerical copy change; they are not tests of a constant coding fraction. Joint histories were reconstructed by lexicographic stepwise Sankoff parsimony on states constrained to 0≤C≤D: total absolute DNA change was minimized first and coding change second. For unambiguous DNA-gain branches, all optimal histories were used to distinguish no coding gain, a smaller coding gain, an equal coding gain and mixed/uncertain states. Event fractions were bootstrapped by neighbourhood. Historical source-table category names are retained with these definitions.

Independent examples were selected with thresholds frozen before final NLR classification. DH06 was excluded; a candidate required at least eight evaluable genomes, DNA range ≥2, at least two genomes with a strict DNA–coding gap ≥2, replicated inclusive gaps and baseline genomes, inclusive range below DNA range, no more than 5% technically uncertain or multi-neighbourhood loci, and at least two explicitly disrupted non-intact copies. We retained at most five cases on distinct chromosomes and, where reconstructed, distinct gain branches. For those cases and the held-out DH06 calibration locus, the complete recovery pipeline was repeated with 25-kb rather than 12-kb flanks to test boundary sensitivity. Case selection was not altered after inspecting these wider-window results. A separate evidence inventory recorded whether each case had same-accession raw long reads, an independent same-accession reassembly, breakpoint reconstruction and copy-specific lesion validation; these criteria graded mechanism evidence but did not redefine the frozen copy–coding patterns.

### Orthogonal high-confidence copy–coding tests

The complete-motif analysis retained DNA-NLR calls whose motif architecture contained both NB-ARC and LRR signatures. The most restrictive complete_unique_clean_n01 subset further required a unique neighbourhood assignment, no interval-boundary flag and an N fraction ≤0.01 in the extracted locus window. Strict and inclusive coding calls, cell evaluability, model formulae, branch-reconstruction rules and all thresholds were inherited unchanged from the primary analysis. Fixed-effect models, leave-one-genome-out fits, phylogenetic independent contrasts, range tests and joint DNA-gain reconstructions were rerun on each frozen subset. The same selected-case rule was applied independently to each subset, with a maximum of five loci.

### Independent *Arabidopsis* neighbourhood replication

The independent panel used the graph-defined NLR neighbourhoods, assemblies and curated annotations reported for 17 *A. thaliana* accessions (Teasdale et al. 2025). We retained NLR_neighbourhood_ID as the homology unit and performed no cross-species re-clustering. The released BED contained 125 unique identifiers, whereas official Supplementary Table S4 contained 123. The published Methods state that two neighbourhoods containing only one incomplete NLR fragment in one accession were removed. We applied that rule by requiring exactly one Curated_type=nlr record, in one accession, annotated as pseudogene or pseudogenic_region; this identified chr3_nbh15 and chr4_nbh02 and yielded the published total of 121. The primary analysis used this reconstructed 121-ID set. The released 125-ID set and the 118-ID subset containing at least one Curated_type=nlr, Gene_status=gene model were analysed as denominator sensitivities. DNA calls containing both NB-ARC and LRR were designated complete motif. A strict DNA–coding link required a unique intact model and reciprocal call–model overlap ≥0.50; the inclusive link required an intact model overlapping at least 0.25 of the shorter feature. Binomial linked-coding-fraction models and Poisson log-link count models included neighbourhood and accession fixed effects with neighbourhood-clustered uncertainty. The count model used log DNA, restricted inference to DNA-positive cells and retained zero coding counts. Independently curated gene totals may exceed DNA-call totals, so they were modelled as counts, with linked-count and curated-only denominator sensitivities reported separately. In the frozen published-121 set, both models were rerun after retaining only DNA-variable neighbourhoods and after excluding the five neighbourhoods with the greatest panel-wide DNA-copy totals. Every neighbourhood and every accession were then omitted in turn. Curated-status sensitivities contrasted (i) gene against protopseudogene, pseudogene and pseudogenic_region; (ii) gene against protopseudogene and pseudogene after excluding pseudogenic_region; and (iii) explicit gene against explicit pseudogene. Range attenuation was tested independently for each definition with ties excluded from the exact sign test. The *Capsicum* and published-121 strict-fraction log odds ratios were compared by their difference under independent-system standard errors; any pooled estimate is descriptive two-system concordance, not a plant-wide meta-analysis.

### Matched RNA-seq and Iso-Seq transcript evidence

Ten paired-end RNA-seq libraries from PRJNA962192 were mapped only to their matched G1 or Andean T2T genome with HISAT2 v2.2.1 (Kim et al. 2019). Overlapping fragments were assigned to DNA-NLR intervals with featureCounts v2.1.1 using -O --fraction (Liao et al. 2014).

Strict transcript support required either ≥5 MAPQ≥20 fragments and CPM ≥0.1 in one tissue, or at least two tissues each with ≥2 fragments and CPM ≥0.05. An inclusive call required ≥2 overlapping MAPQ≥0 fragments in at least one tissue. The four frozen coding classes were strict intact, inclusive only, explicitly disrupted and no intact coding model. Wilson intervals were reported for every class. Cluster-robust logistic models tested locus-level support while adjusting for log locus length, log1p neighbourhood DNA copy number and genome; locus-by-tissue models additionally adjusted for tissue and clustered observations by locus.

For *Arabidopsis*, accession-matched Iso-Seq reads from PRJEB91362 were aligned to their corresponding assembly with minimap2 v2.31-r1302 in splice-aware long-read mode, using annotated splice junctions. MAPQ ≥20 primary alignments were processed with SAMtools v1.21 and intersected with DNA-NLR calls. Read names, splice-junction CIGAR operations and all simultaneously overlapped DNA calls were tracked explicitly; a locus-unique read overlapped exactly one DNA-NLR call in that accession. Support was reported at thresholds of at least one and two locus-unique reads and separately for splice-junction-bearing reads. TAMA-collapsed transcript structures and TransDecoder complete-ORF models from the same accession-matched Iso-Seq release were separately intersected with DNA calls, with locus-unique, multi-exon and ≥80%-span states retained separately. Accession at6137 was excluded because no matched public Iso-Seq run was available. Logistic overlap models adjusted for locus length, neighbourhood DNA copy number and accession and clustered by neighbourhood. Transcript evidence was not used to redefine coding state. These Iso-Seq resources informed the original annotation study and were not a held-out validation set. Complete-ORF status describes the transcript prediction and does not require a complete NLR protein architecture.

### Outgroup and repeat context

Outgroup polarization, local trees and matched NLR-free repeat comparisons characterized the structural background of neighbourhood turnover. Local alignments defined homologous intervals, while a common-library Dfam sensitivity assessed repeat-annotation differences. Software, thresholds, matched-control construction and sequence-selection rules are provided in Supplementary Methods, under “Detailed supporting analyses”.

### DH06 reciprocal families and coding-model rescue

DH06 was bounded by the conserved anchor shell of PNOU_DH06_N0OG0010401_N0OG0019490_001. G1 order defined the F392–F402 family axis. Assignments required reciprocal positional compatibility plus local nucleotide or protein evidence; ambiguous and non-reciprocal DNA-NLR loci remained visible but outside family copy counts. Twenty-six reciprocal-family loci without reliable models in G1, Zhangshugang, PI 632928 or Grif1614 were analysed by family-matched miniprot projection, AUGUSTUS v3.5.0 with tomato parameters in 12-kb flanks, HMMER v3.4 search against PF00931 and NLRtracker (Stanke et al. 2006; Eddy 2011). Predicted features and coding sequences were normalized with gffread v0.12.9 (Pertea and Pertea 2020). Final states prioritized family-matched coverage and identity, frameshifts and in-frame stops over domain presence alone. For literature-coordinate reconciliation, the Liu et al. interval was converted from 1-based inclusive to the same 0-based half-open Zhangshugang coordinate system used by CaNLOG; overlap was calculated separately for the broad anchor shell, F392–F402 core, primary Grif1614 homology segments and nearest CROSSCOPY breakpoint-flanking projections. Primary PAF segments required tp:P, MAPQ≥20 and alignment-block length≥5 kb; their union is reported as a discontinuous homology footprint, not a projected continuous duplication.

### Block duplication, structural breakpoints and copy-specific lesion phasing

The two Grif1614 DH06 copies were self-aligned with minimap2 and reduced to a monotonic same-orientation chain. Exact alignment operations defined large copy-specific sequences, which were rerun through the targeted repeat workflow. Reciprocal-family pairs were assessed in 100-kb windows. Coding lesions were called from amino-acid-aware miniprot alignments and compared by affected codon, genomic coordinate and the physical copy offset. For the three disrupted pairs, the previously projected genomic span plus 10 kb on each side was extracted from each copy and aligned with minimap2 using -x asm5 -c --eqx --cs=long --secondary=no. Non-overlapping, collinear primary segments were combined. Identity was calculated from A/C/G/T matches divided by aligned columns including inserted and deleted bases, separately for each genomic span and flank; unaligned sequence was reported through query coverage. Genomic spans include introns. This comparison was descriptive and did not test gene conversion or date lesion origin (Supplementary Table 26).

Grif1614 HiFi reads were mapped competitively to the complete genome. Structural edges required primary reads spanning both flanks without a discordant large indel. For copy-specific lesion tests, a 201-bp reference context centred on each of 18 lesion entries was searched exactly in target-region HiFi reads. A read supported a physical copy only when the same read both contained the exact lesion context and had a primary Chr06 alignment extending ≥100 bp on each side of that lesion at MAPQ ≥20. Strict support additionally required the primary alignment score to exceed the best competing Chr06 alignment by ≥100. Locus-level phasing required intersection of read IDs across every shared lesion in that locus.

Andean continuity was tested by competitive whole-genome mapping of SRR24319436. Target coverage, depth in 5-kb bins and primary reads spanning both inner-anchor edges were calculated independently of the NLR caller.

### Population context of the DH06-derived module

Junction calling and local-assembly procedures for the 38-accession *C. pubescens* comparison are described in Supplementary Methods, “Population context of the DH06-derived module”.

### Neighbourhood influence and high-copy recovery diagnostics

Each of the 173 pepper neighbourhoods was omitted in turn under the unchanged primary Poisson model and all three coding definitions. We retained coding-zero observations in DNA-positive cells, neighbourhood/accession fixed effects and neighbourhood-clustered uncertainty; the three complete-data rechecks reproduced the published primary estimates. The original 1,023 held-out complete models were partitioned into DNA-copy bins 1, 2, 3–4, 5–15, 16–22 and >22, using the existing 15/22 sensitivity cutoffs. Bin-level Wilson intervals are descriptive locus-level intervals, not cluster-adjusted comparisons. Binomial logit models adjusted for genome and log locus length, used 5–15 as the reference bin, and estimated neighbourhood-clustered intervals. All coefficients and exploratory unadjusted P values were retained (Supplementary Tables 11–12).

### Interpretation of coding states and graphical summaries

Coding-state labels describe sequence-based predictions; receptor activity was not measured. Figure 1a illustrates the positional comparison unit. The curves in Fig. 2a use the full-range count model; their bands transform confidence limits for the count elasticity, whereas the accompanying histogram describes the observed DNA-count distribution. Raw ranges, phylogenetic contrasts and reconstructed branch increments summarize numerical copy differences separately from the proportionality test. The 25-kb flank rerun assesses extraction-window robustness. Structural support for the selected cases was recorded separately: same-accession long reads and reassemblies were unavailable for PI 632928, and the recurrent chromosome 9 pattern lacked copy-specific breakpoint reconstruction. Family-level curves in Fig. 4 summarize positional correspondence, and the Andean coverage panel is schematic. Figure 5e displays observed edge-count minima and maxima; individual reads and structural edges are technical observations. *Arabidopsis* Iso-Seq contributed to the source annotations, so the transcript comparisons describe related evidence rather than independent validation. Transcript detection, predicted ORF completeness and NLR protein architecture were evaluated as separate properties.

### Statistical analysis and figure production

Genomic interval arithmetic used BEDTools v2.31.1 (Quinlan and Hall 2010). Primary count and fraction inference used neighbourhood-clustered standard errors and two-sided 95% intervals.

Count-model tests were against β=1; fraction-model tests were against a log odds ratio of zero. Sensitivity fits are reported in full without significance filtering and their exploratory P values are unadjusted. Exact directional range sign tests and historical bootstrap summaries are retained as descriptive supporting analyses. Group comparisons used 100,000 label permutations, and the nine-point Spearman test enumerated all 9! permutations. Multiple related event tests were adjusted by Benjamini–Hochberg where applicable (Benjamini and Hochberg 1995). Analyses used Python, R, NumPy, pandas, statsmodels and SciPy; figure production used Matplotlib. Figures were exported as editable SVG, vector PDF, 300-dpi PNG and 600-dpi LZW-compressed TIFF (Harris et al. 2020; Virtanen et al. 2020; Hunter 2007), with tab-delimited source data for quantitative panels.

## Data availability

All assemblies, annotations, variants and sequencing reads analysed here were previously public. Assembly accessions, BioSamples, BioProjects, sequence roles and checksums are listed in Supplementary Table 1. Key long-read datasets are Grif1614 SRR23352910 (PRJNA801499) and Andean SRR24319436 (PRJNA962192). The 38 population run accessions derive from PRJNA801499 and are listed in Supplementary Table 27; per-chromosome public-VCF URLs, byte lengths, server modification times and ETags are included in the source-data bundle. The independent *Arabidopsis* genomes, annotations and neighbourhood calls were obtained from Zenodo record 17945820, and accession-matched Iso-Seq runs derive from PRJEB91362 (Teasdale et al. 2025). Matched G1 and Andean RNA-seq runs SRR28789546–SRR28789555 derive from PRJNA962192 and are listed in Supplementary Table 28. The external pepper call-set comparison used Dong et al. Supplementary Table S5 (Dong et al. 2026).

Processed data and source data underlying the reported tables and figures will be deposited together with the analysis code in a Zenodo archive. The repository link will be added in a subsequent version of this preprint.

## Code availability

Project-specific scripts, configuration files and software-version information required to reproduce the analyses, tables and figures will be deposited in the same Zenodo archive described above. The repository link will be added in a subsequent version of this preprint.

## Acknowledgements

The authors thank the Pepper Research Group in the College of Horticulture for providing computational resources and access to the computing server used in this study.

## Author contributions

Liming Xiong conceived and designed the computational framework of the study, performed and supervised the bioinformatic analyses, interpreted the results, prepared the figures, and drafted the manuscript. Feng Liu provided primary supervision and guidance throughout the study, contributed to the interpretation of the results, and critically revised the manuscript. Xuexiao Zou provided additional supervision and scientific guidance, contributed to the interpretation and discussion of the results, and critically reviewed and revised the manuscript. All authors read and approved the final manuscript.

