## Supplementary Information for "NLR DNA copy-number expansion outpaces accumulation of predicted coding-intact copies"

### Supplementary Methods

#### Evidence hierarchy and evaluability

CaNLOG retains positional homology, DNA-NLR presence, predicted coding integrity and transcript support as separate variables. A PNOU×genome cell is assigned a biological zero only when its anchor shell is uniquely delimited, continuously assembled and fully scanned without a DNA-NLR call. Broken or ambiguous shells remain `NOT_EVALUABLE`.

#### Complete-motif and technically clean subsets

The complete-motif subset requires both NB-ARC and LRR motif signatures. The complete/unique/clean subset additionally requires unique PNOU membership, no interval-boundary flag and N fraction $\leq 0.01$. Primary strict/inclusive coding definitions, model formulae, phylogenetic contrasts and branch-reconstruction rules are unchanged. Fixed-effect, leave-one-genome-out, range, PIC and DNA-gain analyses were rerun independently in each frozen subset.

#### Direct-count, recovery-bias and sampling sensitivities

The primary estimand is the conditional count elasticity from a Poisson log-link model, log E(C)=neighbourhood effect+accession effect+$\beta$ log(D). DNA-zero cells are outside this model, while coding-zero cells are retained. At $\beta=1$ the expected coding fraction is constant conditional on the two fixed effects. Two-sided Wald intervals and tests against one use neighbourhood-clustered sandwich covariance. All model estimates, convergence flags, sample domains and two-sided and directional probabilities are supplied in Supplementary Table 10. Sensitivities include fully evaluable neighbourhoods, exact four-anchor cells, DNA$\leq$15 and DNA$\leq$22 cells, species-balanced panels, omission of each accession, omission of DH06, complete-motif, unique-membership and N-content restrictions, and coding thresholds. These are sensitivity comparisons, not a collection of independent replications.

Earlier log1p-on-log1p slopes and positive-count hurdle fits are retained in the code/data archive as descriptive transformed-count models. The log1p slope at constant coding fraction need not be one: doubling DNA from 2 to 4 and coding count from 1 to 2 gives slope log(3/2)/log(5/3)=0.794 despite a constant fraction of 0.5. Raw ranges, raw PIC slopes and equal branch increments likewise test equal numerical change rather than a general constant fraction. Their observed values remain useful descriptions and are not the primary proportionality tests.

For calibration, 1,023 complete published models with unique PNOU membership were hidden from all recovery routes and stratified by DNA-copy count. Genome-adjusted logistic models tested copy-dependent recovery. Two 1,000-replicate binomial simulations tested the coding-fraction model under copy-independent coding probability: one retained fitted PNOU and genome effects and the other imposed empirical copy-bin recovery. These simulations calibrate coding-fraction odds ratios, not log1p count elasticities. Add-one probabilities compare the simulated and observed fraction coefficients.

#### Assigned and unassigned locus universe

All 10,792 DNA-NLR loci were retained and classified as anchor-assigned or unassigned. Complete-motif status, internal-stop flag and primary-sequence role were compared by Fisher's exact tests. Locus length, motif count, distance to sequence edge and distance to the nearest N-run were compared by Mann–Whitney tests with rank-biserial effects. The inferential claim was restricted to the 6,555 anchor-assigned loci; unassigned loci were not treated as biological absences.

### Independent *Arabidopsis* replication

The published graph-defined `NLR_neighbourhood_ID` was retained as the homology unit for 17 *A. thaliana* accessions. The released BED contains 125 identifiers and official Supplementary Table S4 contains 123. Following the article's exclusion rule, `chr3_nbh15` and `chr4_nbh02`—each represented by one curated NLR pseudogene in one accession—were removed from Table S4 to reconstruct the published 121-neighbourhood set. The 125-ID released superset and the 118-ID subset containing a curated intact NLR gene were retained as denominator sensitivities. Complete-motif DNA calls contain both NB-ARC and LRR. Canonical coding models require `Curated_type=nlr` and `Gene_status=gene`. Strict linking requires a unique model with reciprocal overlap ≥0.50; inclusive linking requires an intact model covering at least 0.25 of the shorter feature. Coding-fraction models include neighbourhood and accession fixed effects. Poisson log-link models with log DNA estimate count elasticity on DNA-positive cells. Curated gene counts are not required to be a subset of DNA calls; linked-count and curated-status-denominator analyses distinguish these estimands. The full count-relation matrix, including DNA-zero cells, is retained in Supplementary Table 21.

Primary inference used the reconstructed published-121 identifiers. Two subset sensitivities retained only neighbourhoods whose all-DNA copy count varied across accessions or excluded the five neighbourhoods with the greatest total all-DNA copy count across the 17-accession panel. Influence analysis omitted each of the 121 neighbourhoods and each of the 17 accessions in turn and refitted both the strict coding-fraction model and curated-gene direct-count model. Complete estimate ranges, extreme confidence bounds, maximum directional or two-sided $P$ values and convergence states were retained.

Curated-status sensitivity did not silently absorb `protopseudogene`. Three explicit grouped responses compared (i) `gene` with all degraded states (`protopseudogene`, `pseudogene` and `pseudogenic_region`), (ii) `gene` with `protopseudogene` and `pseudogene` after excluding `pseudogenic_region`, and (iii) explicit `gene` with explicit `pseudogene` after excluding both other degraded labels. Each definition was run in the full published-121 set, the DNA-variable subset and the top-five-excluded subset. For range attenuation, the maximum-minus-minimum curated-gene count across accessions was compared with the corresponding denominator range; ties were excluded from an exact one-sided sign test. This range summary was analysed independently of the fixed-effect regression.

The primary strict coding-fraction log odds ratios from *Capsicum* and published-121 *Arabidopsis* were compared as independent-system estimates. Their difference, standard error, 95% CI and two-sided $P$ value were calculated on the log-odds scale; the exponentiated difference is the ratio of odds ratios. Cochran's $Q$ and an inverse-variance fixed-effect pooled estimate are provided only as descriptive two-system concordance. They are not interpreted as a plant-wide meta-analysis.

### DH06 literature-coordinate reconciliation

The Liu *et al.* Zhangshugang interval Chr06:242,250,001–246,150,000 was treated as 1-based inclusive and converted to Chr06:242,250,000–246,150,000 in the 0-based half-open CaNLOG convention before overlap calculations. It was compared separately with the broad Zhangshugang DH06 anchor shell, the F392–F402 reciprocal-family core, the project-local approximate WP1C interval, primary Grif1614-to-Zhangshugang homology segments and the nearest reliable CROSSCOPY breakpoint-flank projections. Grif1614 PAF segments were retained at `tp:P`, MAPQ≥20 and alignment block length≥5 kb. Because these segments are discontinuous, their minimum-to-maximum coordinate span is labelled a homology footprint rather than a continuously projected duplication.

#### Evidence grading for non-DH06 replication cases

The five frozen non-DH06 cases were retained as independent chromosomal copy–coding patterns. The 25-kb rerun was a flank-width sensitivity of the same assemblies and was not counted as new raw-read or independent-assembly evidence. A focused inventory recorded, for DH01 PI 632928 and the DH09 *C. annuum* pattern, whether current inputs contained same-accession raw long reads, an independent same-accession reassembly, breakpoint-level reconstruction or copy-specific raw-read lesion validation. Multiple independently assembled *C. annuum* materials were considered recurrence evidence for DH09 architecture, but not a same-accession reassembly or a DH06-equivalent breakpoint mechanism.

#### Pepper RNA-seq validation

Ten PRJNA962192 paired-end libraries were aligned only to their matched G1 or Andean T2T genome with HISAT2 v2.2.1. featureCounts v2.1.1 used `-O --fraction` to retain fractional evidence for overlapping NLR loci. Strict support requires either ≥5 MAPQ≥20 fragments and CPM ≥0.1 in one tissue, or at least two tissues each with ≥2 fragments and CPM ≥0.05. Inclusive support requires ≥2 MAPQ≥0 overlapping fragments in at least one tissue.

#### *Arabidopsis* Iso-Seq validation

PRJEB91362 Iso-Seq reads were aligned to the matched accession assembly with minimap2 v2.31-r1302 in splice-aware long-read mode using annotated junctions. MAPQ ≥20 primary alignments were filtered with SAMtools v1.21. A read was locus unique when it overlapped exactly one DNA-NLR call in that accession. CIGAR `N` operations identified splice-junction-bearing reads; reference-span ≥80% and aligned-call-bases ≥50% were retained separately. Accession-aware transcript identifiers prevented models with reused names from collapsing across assemblies. TAMA-collapsed structures and TransDecoder models marked `ORF type:complete` were intersected independently with the DNA calls, recording locus uniqueness, multi-exon structure and ≥80% call span. Logistic models for ≥2 unique reads, unique splice support, unique TAMA structure and unique complete-ORF structure adjusted for log locus length, log1p neighbourhood DNA-copy number and accession and used neighbourhood-clustered standard errors. at6137 was excluded because no matched public Iso-Seq run was present in the frozen panel.

Full threshold definitions, software versions, row-level results and accession accounting are retained in Supplementary Tables 10, 13–15, 20–21, 24, 26 and 28–43 and the source-data archive.

#### DH06 paired sequence context

The existing projected genomic span of each disrupted F396, F397 and F401 copy was extracted with 10-kb flanks from the Grif1614 interval FASTA. Minimap2 used `-x asm5 -t 1 -c --eqx --cs=long --secondary=no`. Non-overlapping collinear primary segments were combined, with query and target coordinates checked for monotonicity. For each span and flank, matches, substitutions, inserted/deleted bases, query coverage and aligned identity were retained. Identity excludes unaligned sequence and includes indel columns in its denominator. The spans include introns and are not CDS-only alignments. F396 has two primary segments separated by a local difference; both are included. The six extracted FASTAs, PAFs, extraction manifest and comparison script are provided for reproduction. This descriptive comparison assesses the sequence context shared by damaged pairs; it does not date damage or formally infer gene conversion.

The Iso-Seq resource contributed to the original Arabidopsis curated annotations. It is related transcript evidence, not an independent held-out validation; a complete transcript ORF does not establish complete NLR protein architecture.

### Detailed supporting analyses

#### Controlled perturbation benchmark

The benchmark truth set comprised high-confidence projected anchor shells in chromosome-resolved G1 and Grif1614. Units required high projection confidence, a complete/partial DNA-NLR state or an evaluable no-call, and at least three mapped anchors. This produced 274 genome-specific units: 255 DNA-NLR-present units and 19 evaluable no-call units. Four representations were compared: CaNLOG non-NLR anchor shells with DNA-defined target state; annotation-dependent anchor synteny requiring a protein NLR; protein HOGs usable only when positionally pure; and single-linkage NB-ARC sequence clusters at ≥70% identity over at least 30 aligned residues. Sequence-cluster thresholds of 60%, 70% and 80% were retained as sensitivity results.

For each of 200 fixed-seed replicates per condition, 10%, 20% or 40% of NLR annotations, all annotations, or usable gene models were removed; contig boundaries were simulated at 100-, 25- or 5-Mb spacing with random chromosome offsets; and 5%, 10% or 20% of positionally unique protein-HOG calls were swapped between paralogous units. Recovery, specificity, precision, called-state accuracy, false-loss rate and unresolved fraction were computed against the unperturbed high-confidence units. A CaNLOG shell crossing a simulated boundary was unresolved. The benchmark is a recovery stress test conditional on its truth-set construction, not an experimental estimate of genome-wide method accuracy.

#### Candidate disclosure, outgroup polarization and local trees

The complete state-contrast universe contained 30 PNOUs with both G1–Andean state contrast and paired interval/repeat measurements. Nine initial structural priorities and 15 full-screen extensions were selected post hoc and labelled exploratory. Ranking variables, within-wave ranks, six non-selected cases and a grid of occupancy, length-ratio and repeat-difference thresholds are provided in Supplementary Table 22.

For followed PNOUs, homologous intervals were delimited in tomato, potato, *C. chacoense* and *C. galapagoense* using compatible anchor pairs. Protein evidence, NLR-Annotator calls and local nucleotide alignment classified each interval as present, absent-not-detected or not evaluable. Failed anchors were never coded absent. Directional labels summarize the most parsimonious outgroup pattern and not an exact mutational mechanism. NB-ARC or full-length protein sequences were aligned with MAFFT and exploratory trees inferred with FastTree v2.2.0 (Price et al. 2009); short, invariant or sparsely sampled alignments were labelled low information.

#### Local alignment, repeat annotation and matched controls

Local nucleotide alignment used minimap2 v2.31-r1302 with `-x asm20 -c --eqx --cs=long --secondary=yes -N 20` (Li 2018). Coverage, identity, orientation and discontinuities were derived from PAF, CIGAR and `cs` operations. Published genome-wide repeat tracks provided the primary screen. Target intervals were independently rerun with RepeatMasker v4.2.4, RMBlast 2.17.1+ and the same *C. annuum*-clade Dfam 4.0 library (NCBI taxon 4072 with ancestors) (Nishimura 2000; Storer et al. 2021). Between-source direction concordance and Spearman correlation were reported by repeat class.

For each of the 78 primary G1–Andean PNOUs, a unique non-NLR control was selected from four consecutive conserved anchors with no Stage-4 DNA-NLR call in either genome. Controls were matched on G1 and Andean interval lengths and chromosome where possible. Repeat effects were evaluated as between-genome percentage-point differences and as paired PNOU-minus-control differences.

#### Population context of the DH06-derived module

Junction-edge evidence and focused local assemblies classified 38 public C. pubescens accessions, retaining unresolved samples separately. Local SNP comparisons were exploratory; genome-wide structure and Firth regression assessed geographic enrichment conditional on measured SNP structure. Detailed calling, assembly, filtering and regression procedures are retained in the corresponding existing supplementary methods.

#### Population junction genotyping and callability

Five exact 51-mer markers per edge first detected the alternative CROSSCOPY junction. Reference callability was then evaluated with 50 unique markers in each flanking reference interval, estimated haploid depth and stable control loci. An alternative was supported by both exact edges or by a continuous, 100%-identity focused assembly spanning the locus. A noncarrier required global depth ≥5×, at least 10 nonzero reference markers per flank and a median positive count ≥2 on both flanks. All other alternative-negative samples were unresolved. Target-matching read pairs were assembled with SPAdes v4.3.0 at k-mers 21, 33, 55, 77, 99 and 127 (Bankevich et al. 2012), and candidate scaffolds were evaluated with nucleotide alignment and miniprot. Legacy H1–H5 labels summarize combinations in this same junction-event matrix; they are descriptive profiles, not independently phased haplotypes, and were not used as predictors in the callability or Firth analyses.

#### Local SNP context of PI 585272

Local small-variant context was evaluated across Zhangshugang Chr06:244,896,574–250,579,355, comprising the anchor-defined DH06 interval and 1-Mb flanks. One 50-kb segment was surveyed within each of 29 consecutive 200-kb bins (1.45 Mb in total). Within the 38-sample VCF, DP <3 or GQ <10 genotypes were set to missing and sites were retained at missingness ≤20% and MAF ≥5%, leaving 1,047 common SNPs. Physical thinning at 1-kb spacing retained 185 SNPs for local PCA and pairwise standardized distances; 180 remained polymorphic in the standardized-distance calculation. All 1,047 SNPs were retained for per-segment information, missingness and mean absolute genotype-distance contrasts.

Resolved comparator groups comprised 19 Guatemalan alternative carriers, four Guatemalan callable noncarriers and seven Ecuadorian callable noncarriers. The three unresolved accessions remained visible in the PCA and complete distance matrix but were never included in a negative comparator group. For each surveyed segment, the plotted contrast was the mean distance from PI 585272 to Guatemalan alternative carriers minus its mean distance to Ecuadorian callable noncarriers; negative values indicated greater similarity to the Guatemalan carrier group. The cross-copy homology zone was defined by the nearest reliable flanking alignments projected to Zhangshugang, not by the local SNP pattern itself. This analysis was exploratory and did not assign a unique historical mechanism.

#### Genome-wide population structure and Firth regression

The public Zhangshugang VCF was processed for all 38 accessions chromosome by chromosome with BCFtools (Danecek et al. 2021). Chr06 was excluded a priori to avoid proximal contamination from DH06. PASS biallelic SNPs were retained after setting genotypes with DP <3 or GQ <10 to missing, then requiring site missingness ≤20% and MAF ≥5%. Variants from the remaining 11 chromosomes were concatenated and pruned in 100-kb windows at pairwise $r^2$

≤0.20. If more than 50,000 pruned variants remained, a fixed-seed sample of 50,000 was used. PLINK 2 generated the complete 37-axis non-zero eigenspectrum and a KING kinship matrix; because the cohort contained fewer than 50 accessions, allele frequencies estimated from this post-QC fixed-seed cohort were supplied explicitly for variance standardization. The first ten PCs were retained in source data and up to four were used for regression adjustment (Chang et al. 2015). Unknown origins were retained as a separate display class but excluded from origin-effect models; unresolved local genotypes were also excluded from binary models.

Firth bias-reduced logistic regression modelled the alternative-supported state against Guatemala origin with zero, two or four standardized PCs (Firth 1993). Penalized likelihood-ratio $P$ values were obtained by fixing the origin coefficient to zero within the same full model and maximizing over nuisance coefficients; 95% confidence intervals were obtained from the corresponding penalized profile likelihood. The adjusted-score solution was required to agree with direct numerical maximization of the Jeffreys-penalized likelihood. A profile bound that could not be bracketed was retained as unbounded rather than truncated to a finite value. Nearest-neighbour event concordance was summarized from the KING matrix. These models test whether geographic enrichment remains after measured SNP structure; they do not estimate adaptation or causal geographic effects.

#### Pepper influence and high-copy diagnostics

We used the frozen 1,741 evaluable pepper cells and the existing Poisson log-link estimator with log DNA, neighbourhood and accession fixed effects and neighbourhood-clustered covariance. DNA-zero cells remained outside the log-DNA estimand, while coding-zero cells were retained. For strict, inclusive and AUGUSTUS/domain definitions, 173 neighbourhood omissions and one complete-data recheck were fitted (522 total fits). Complete-data estimates matched the released primary coefficients to $<10^{-7}$. No significance-based selection was applied. We report full β ranges, largest confidence upper bounds, maximum two-sided P against β=1, convergence and the largest absolute influence.

Recovery diagnostics reused the 1,023 held-out complete models and unchanged locus recovery calls. Bins 1, 2, 3–4, 5–15, 16–22 and >22 refine the original bins using the existing 15/22 sensitivity boundaries. Wilson intervals describe binomial proportions at the locus level and do not account for clustering. Complementary binomial logit diagnostics use these bins, genome and log locus length as predictors, reference D=5–15, and PNOU-clustered covariance. All bin contrasts, including nonsignificant results, are retained with exploratory unadjusted P values. Calibration sample coverage is explicitly reported by neighbourhood and genome.

### Supplementary Results

#### Population context of the DH06-derived cross-copy module

Exact junction edges and focused assemblies identified a DH06-derived structural module in 38 public C. pubescens accessions. Twenty carried the alternative module, 15 were callable noncarriers and three remained locally unresolved. The junction intersects the final coding exon of the divergent NLR-like model Cpu06g32690; five representative carrier assemblies shared replacement of the reference C terminus followed by a premature stop. The alternative can coexist with reference-span evidence, identifying a segregating cross-copy structural module rather than a simple biallelic deletion. Population structure, geography-adjusted tests and the exploratory PI 585272 local-haplotype analysis are reported in Supplementary Figs. 6–7 and Supplementary Table 27.

#### Direct coding-copy counts establish sublinear scaling

Among 1,622 DNA-positive pepper cells, strict, inclusive and AUGUSTUS/domain count elasticities were 0.565 (95% clustered CI 0.345–0.785; two-sided P=0.000104), 0.879 (0.779–0.979; P=0.0181) and 0.891 (0.775–1.006; P=0.0628). The last interval includes proportional growth. Strict point estimates across 11 leave-one-accession-out fits ranged from 0.464 to 0.611, with every interval below one; seven species-balanced estimates ranged from 0.584 to 0.650, again with every interval below one. Inclusive intervals were below one in 10/11 accession omissions and 5/7 balanced panels; the corresponding AUGUSTUS numbers were 2/11 and 4/7. Fully evaluable and exact-anchor strict fits retained the relationship. Restricting cells to DNA≤15 or DNA≤22 weakened it to 0.845 (0.607–1.084) and 0.887 (0.683–1.091). Highly expanded cells thus contribute substantially to the full-range association. Removing DH06 retained $\beta$=0.654 (0.463–0.845). Full estimates are reported without significance filtering in Supplementary Table 10.

#### Held-out models and null simulations quantify measured recovery bias

The held-out validation contained 1,023 complete published models with unique PNOU membership. Inclusive recovery was 80.5%, 81.0%, 75.4% and 76.1% in cells with 1, 2, 3–4 and ≥5 DNA copies, while projection-independent AUGUSTUS/domain recovery was 76.6%, 73.4%, 73.2% and 73.7%. Stringent projection was intentionally less sensitive (35.2%, 31.6%, 38.7% and 34.3%) but likewise did not deteriorate with copy number. Copy number was not a significant negative predictor of held-out recovery in genome-adjusted strict or projection-independent models. Under a constant coding-probability null, the simulated coding-fraction odds ratio was centred near one (median 1.001; 95% simulation interval 0.883–1.133); applying the empirical copy-bin recovery profile yielded a median of 0.992 (0.877–1.117). Only 1 of 1,001 rank positions, after add-one correction, was as low as the observed odds ratio of 0.664 in either simulation.

#### Orthogonal sequence definitions preserve the pepper effect

Complete-motif counts gave strict $\beta$=0.591 (95% CI 0.377–0.805) and inclusive $\beta$=0.876 (0.808–0.944). Requiring complete motifs, unique membership and clean sequence gave $\beta$=0.536 (0.314–0.759) and 0.863 (0.801–0.925). The complete-motif strict coding-fraction OR was 0.648 (PNOU-clustered CI 0.527–0.796). Raw-range summaries contained 100 attenuated versus six amplified PNOUs in the complete-motif set, and 98 versus five in the complete/unique/clean set; both yielded 91/108 DNA-gain events with smaller or absent coding gains. These raw summaries remain descriptive; count and fraction inference is reported in Supplementary Tables 9–10.

#### The anchor-evaluable universe is explicit and biologically representative in motif completeness

CaNLOG assigned 6,555 of 10,792 DNA-NLR loci (60.7%) to the 173 positional units and retained 4,237 outside that inference universe. Complete-motif fractions were nearly identical in assigned and unassigned loci (61.37% versus 61.03%; Fisher's exact $P$=0.731), as were primary-chromosome fractions (25.66% versus 26.08%; $P$=0.636). Assigned loci were farther from assembly edges and N-runs, consistent with the availability of an intact anchor shell. The main claim is therefore stated for anchor-evaluable neighbourhoods, while all unassigned loci remain in the catalogue rather than being recoded as absence.

#### An independent plant panel reproduces regression-level sublinear coding-copy scaling

The published 121-neighbourhood set was reconstructed from the 123 official Table S4 identifiers using the original exclusion rule, with the 125-ID released BED and 118-ID intact-gene set retained as sensitivities. Among DNA-positive cells, curated-gene count elasticity was 0.708 (95% clustered CI 0.571–0.844; two-sided P=0.0000280); strict-linked and inclusive-linked elasticities were 0.785 (0.651–0.918) and 0.808 (0.723–0.893). Using complete-motif DNA as predictor gave curated-gene β=0.640 (0.502–0.777). Estimates were effectively unchanged in the released-125 and intact-gene-118 sets. The strict coding-fraction OR was 0.676 (0.510–0.895; P=0.00630).

Among 70 DNA-variable neighbourhoods, curated-gene count elasticity was 0.706 (0.568–0.843). Excluding the five largest DNA-copy neighbourhoods gave β=0.603 (0.418–0.789). Across all 121 leave-one-neighbourhood-out refits, curated-gene elasticities ranged from 0.671 to 0.751 (largest CI upper bound 0.857; largest two-sided P=0.000236); strict-linked elasticities ranged from 0.750 to 0.816 (largest upper bound 0.936; largest P=0.00451). All 17 leave-one-accession-out intervals also remained below one. Full count fits are in Supplementary Table 10 and retained coding-fraction influence results are in Supplementary Table 43.

Curated-status definitions delimited the count association. Curated genes against a denominator of genes plus all degraded states gave β=0.893 (0.810–0.976; two-sided P=0.0112). Excluding pseudogenic_region while retaining protopseudogene gave β=0.944 (0.870–1.018; P=0.137), and an explicit gene-versus-pseudogene denominator gave β=0.935 (0.877–0.992; P=0.0244). In the last definition the top-five-excluded interval included one (0.943, 0.881–1.005). The corresponding binomial fractions are retained in Supplementary Table 43, including the non-significant OR=0.817 (0.628–1.062; P=0.130) when pseudogenic_region is excluded but protopseudogene retained.

Neighbourhood-level range attenuation was not reproduced in *Arabidopsis*. Under the primary strict-link definition, 17 neighbourhoods were attenuated, 42 tied and 11 amplified; the median DNA-minus-coding range was zero and the exact directional sign test gave $P$=0.172. Curated-gene comparisons likewise had median attenuation zero: gene versus all DNA calls, 10/39/21 attenuated/tied/amplified ($P$=0.985); gene versus all degraded states, 9/53/14 ($P$=0.895); exclusion of `pseudogenic_region` with `protopseudogene` retained, 9/54/15 ($P$=0.924); and explicit gene versus pseudogene, 9/84/6 ($P$=0.304). The cross-system result is therefore sublinear scaling in fixed-effect regressions, not replication of every neighbourhood-level summary statistic.

The strict coding-fraction effects had the same direction across the two independently defined systems: OR 0.664 (0.544–0.811) in *Capsicum* and 0.676 (0.510–0.895) in published-121 *Arabidopsis*. The Capsicum-minus-*Arabidopsis* log-OR difference was −0.0177 (95% CI −0.363 to 0.327; $P$=0.920), equivalent to an OR ratio of 0.982 (0.696–1.387); Cochran's $Q$=0.0101 ($P$=0.920). The non-significant difference does not establish equivalence. The inverse-variance estimate, OR 0.668 (0.567–0.786), is reported only as descriptive two-system concordance and not as a plant-wide meta-analysis.

#### DH06 is distinct from the previously reported Zhangshugang Chr06 peak

In Zhangshugang coordinates, the broad CaNLOG anchor shell spans Chr06:245,896,574–249,579,355 (0-based, half-open). It overlaps 253,426 bp at the distal edge of the Liu *et al.* Chr06:242,250,001–246,150,000 population-level introgression interval after coordinate conversion. In contrast, the F392–F402 reciprocal-family core spans Chr06:247,474,211–248,185,910 and does not overlap that interval; its proximal edge is 1,324,211 bp distal to the published endpoint. The nearest reliable CROSSCOPY breakpoint-flank projections span Chr06:247,506,764–248,194,284 and are likewise non-overlapping. Forty qualifying primary PAF segments from the two Grif1614 blocks define a discontinuous Zhangshugang homology footprint from 246,396,649 to 248,930,305, beginning 246,649 bp distal to the published interval. The project-local approximate WP1C interval, Chr06:242,566,000–243,573,000, lies inside the published peak and is separated from the DH06 core. These coordinates establish adjacency of the broad anchor shell but do not equate DH06 with WP1C, fruit size or a confirmed introgression event.

#### Non-DH06 cases are replicated copy–coding patterns

At DH01, PI 632928 carries five DNA copies but one inclusive predicted coding-intact copy against a one-copy state in seven *C. annuum* assemblies; eight explicit non-intact disrupted loci occur across the frozen case calls. The current project inputs contain neither same-accession PI 632928 raw long reads nor an independent PI 632928 reassembly, so this case is retained as a replicated copy–coding pattern rather than a breakpoint-resolved duplication mechanism. At DH09, seven independently assembled *C. annuum* genomes each contain three DNA copies and zero strict predicted coding-intact copies, with 13 explicit non-intact disrupted loci across the frozen calls. This multi-assembly recurrence supports the architecture, while the absence of same-accession raw-read phasing and copy-specific breakpoint reconstruction distinguishes it from the DH06 mechanism. These optional validations would strengthen individual mechanisms but do not alter the genome-wide or replicated-pattern conclusions.

#### Transcript evidence forms a third state layer

In matched G1 and Andean RNA-seq, strict transcript support occurred in 176/194 strict-intact loci, 277/359 inclusive-only loci, 114/181 disrupted loci and 101/304 loci without an intact model. After adjustment for locus length, PNOU DNA copy and genome, the corresponding odds ratios relative to no-intact-model loci were 10.66 (95% CI 6.29–18.06), 4.09 (2.64–6.32) and 2.43 (1.61–3.67). The tissue-resolved model retained the same ordering.

Accession-matched *Arabidopsis* Iso-Seq added structure-level evidence. Locus-unique splice-junction reads supported 1,325/2,105 (62.9%) coding-linked DNA calls and 656/1,316 (49.8%) copy–coding-gap calls; locus-unique complete-ORF transcript models supported 1,449/2,105 (68.8%) and 704/1,316 (53.5%). Once locus length, neighbourhood DNA-copy number and accession were accounted for, no association of complete-ORF support with the coding-link boundary was detected (OR 1.19, 95% CI 0.78–1.82), nor was an association detected for locus-unique splice support (1.02, 0.67–1.56). Thus, intact coding prediction strongly stratifies short-read transcript detection in pepper, whereas full-length transcript structures in *Arabidopsis* occupy both curated coding-linked and copy–coding-gap DNA states. These confidence intervals do not establish equivalence. The Iso-Seq data informed the original annotation study, and complete transcript ORFs need not encode complete NLRs.

#### Shared damaged templates occupy conserved duplicated sequence

The disrupted F396, F397 and F401 pairs shared two, four and three lesion signatures, with zero private assayed lesions. The genomic spans differed by one base in 3,703 bp, two in 3,670 bp and zero in 3,834 bp, respectively. Left and right 10-kb flank identities ranged from 99.850% to 99.960%. Query coverage was 100% for all but the F396 right flank (99.56% across two primary segments); the remaining 44 query bases are outside the identity denominator. F396 strict read phasing establishes the shared damaged state in both physical copies. F397 has broader copy-specific support and F401 retains only event-level evidence. The sequence context is consistent with amplification of damaged templates but does not distinguish lesion inheritance from later gene conversion. Supplementary Table 26 contains the nine regional comparisons, base-level differences, lesion positions, input manifest and six-pair structural context.

#### Curated and DNA counts have distinct measurement domains

Across 2,057 evaluable cells in the published Arabidopsis set, 437 had both counts zero, 96 had curated genes but no DNA call, 1,417 had positive DNA with curated counts at most DNA counts, and 107 had curated counts exceeding positive DNA counts. All 533 DNA-zero cells remain in the input release. The log-DNA model includes 1,524 positive-DNA cells and 112 neighbourhoods. Curated-gene models measure annotation totals conditional on DNA calls; linked-coding models measure the subset assigned to those calls. Supplementary Table 21 records every relation without silently capping curated counts.

#### Pepper neighbourhood influence and high-copy recovery

All 173 leave-one-neighbourhood-out strict fits retained count elasticities below one with 95% intervals excluding one: β ranged from 0.529543 to 0.653881, the largest upper confidence limit was 0.845218, and the maximum two-sided P against $\beta=1$ was 0.000764086. The corresponding inclusive range was 0.854860–0.898501, with maximum upper confidence limit 0.996189 and maximum P=0.0421449; all 173 intervals excluded one. All fits converged. The largest strict shift followed omission of DH06 (absolute β change 0.0888235). Thus, no single neighbourhood was necessary for the strict or inclusive full-range association. This is distinct from robustness to removing the high-copy range across many neighbourhoods. The AUGUSTUS/domain route gave β=0.864976–0.914240, maximum upper limit 1.023446 and maximum P=0.123756; only 4/173 intervals excluded one (Supplementary Table 12).

The 1,622 DNA-positive cells span D=1–108. Counts in bins 1, 2, 3–4, 5–15, 16–22 and >22 are 698, 265, 274, 307, 35 and 43, respectively. The highest bin spans 17 neighbourhoods across all 11 genomes. Its held-out complete-model calibration comprises 248 loci in 12 neighbourhoods and five genomes, rather than representing every high-copy cell. Strict recovery was 144/357 (40.3%; descriptive Wilson 95% CI 35.4–45.5%) at D=5–15, 22/69 (31.9%; 22.1–43.6%) at D=16–22 and 65/248 (26.2%; 21.1–32.0%) at D>22. Inclusive recovery in these bins was 79.6%, 69.6% and 73.0%; AUGUSTUS/domain recovery was 77.6%, 69.6% and 69.4%, respectively. The >22-versus-5–15 strict recovery contrast adjusted for genome and log locus length gave OR=0.565763 (PNOU-clustered 95% CI 0.290499–1.101856; exploratory unadjusted P=0.093981). Inclusive and AUGUSTUS/domain ORs were 0.799834 (0.438042–1.460440; P=0.467184) and 0.709035 (0.382257–1.315162; P=0.275341). All bin-level counts, all adjusted contrasts and the original 1,023 locus records are released (Supplementary Table 11 and source data).

Finer bins therefore expose recovery heterogeneity that the earlier ≥5 bin pooled together. The imprecise adjusted comparisons neither establish a copy-number effect on recovery nor rule out a biologically meaningful detection deficit in the tail. The original coarse-bin null simulation is not a calibration that incorporates this finer high-tail pattern; it must not be read as excluding all recovery bias. These diagnostics leave the frozen primary coefficients unchanged and sharpen the boundary of their biological interpretation.

### Supplementary figures and legends

**Supplementary Figure 1 | Catalogue completeness, anchor projection and external call-set concordance**

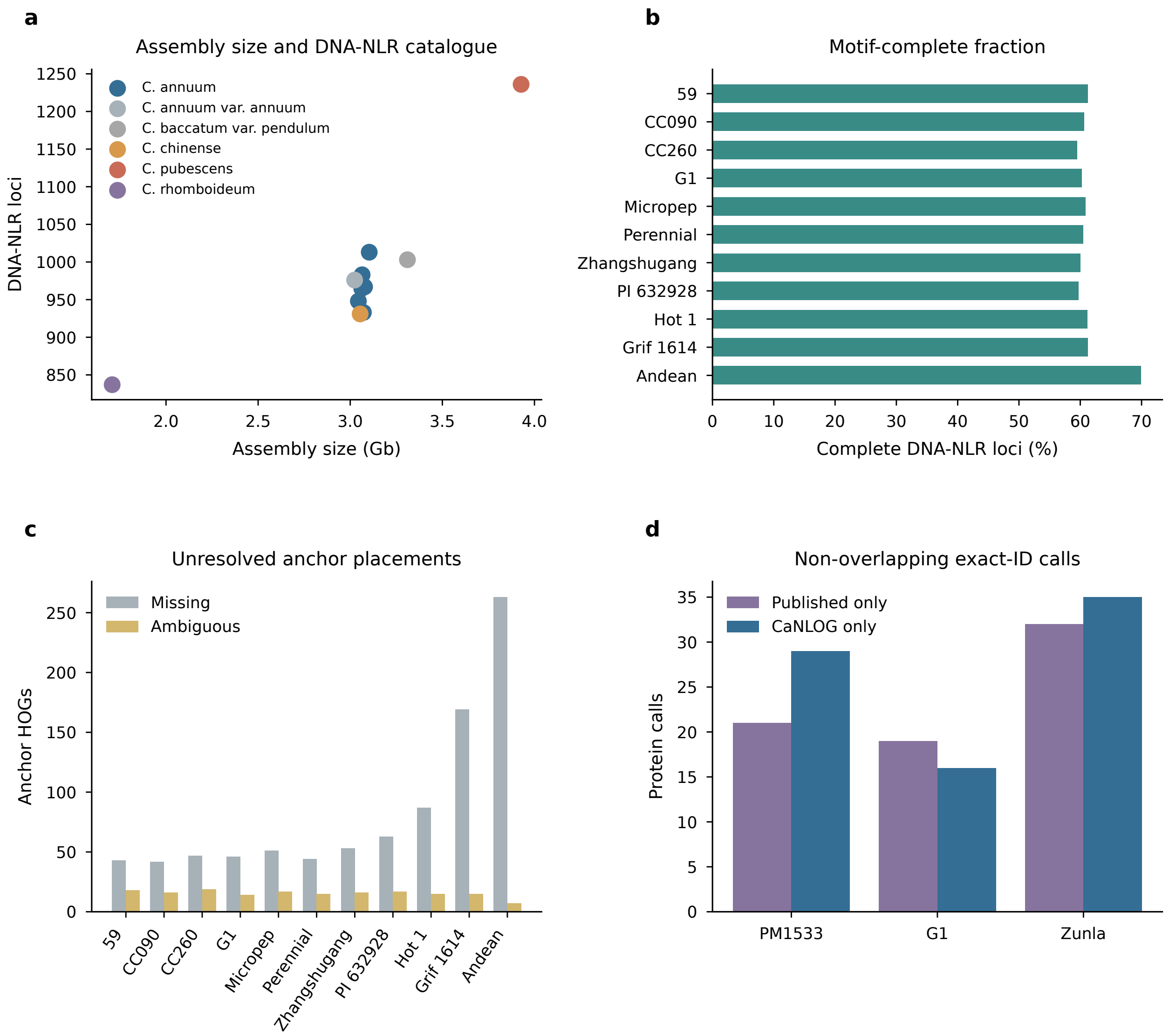


**a**, Assembly size and number of DNA-defined NLR loci in the 11 structural-genome assemblies. **b**, Fraction of DNA-NLR loci with complete motif architecture. **c**, Missing and ambiguous placements after projection of the 1,613 stable anchor HOGs. **d**, Non-overlapping calls in the exact-protein-ID comparison with the 2026 pepper pan-NLRome. The two call sets use overlapping public annotations and are therefore interpreted as methodological concordance rather than independent biological validation.

### Supplementary Figure 2 | Controlled-perturbation recovery benchmark

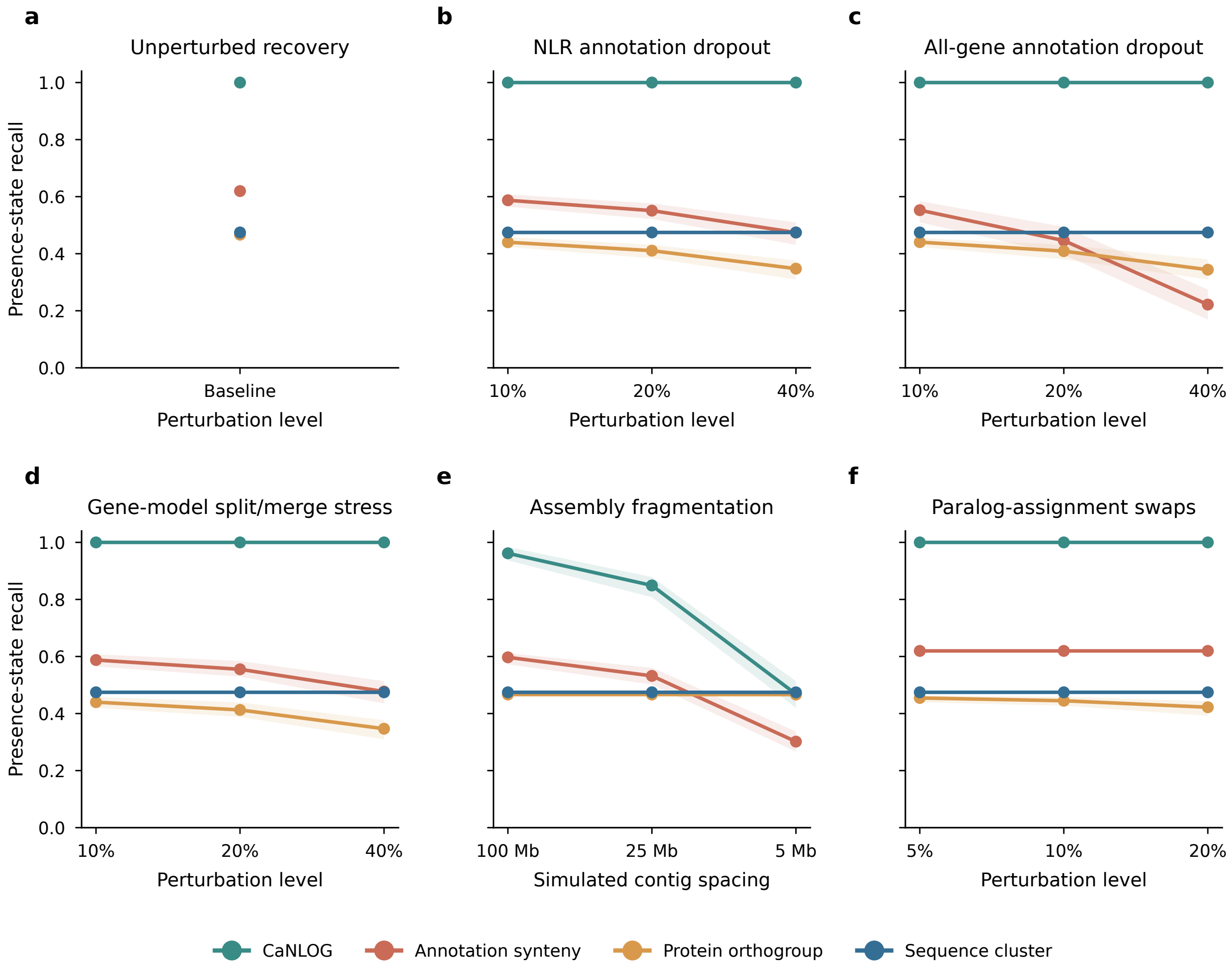


Presence-state recall for CaNLOG, conventional annotation-dependent synteny, protein orthogroups and NB-ARC sequence clusters under **a**, no perturbation; **b**, NLR-annotation dropout; **c**, all-gene annotation dropout; **d**, gene-model split/merge stress; **e**, simulated assembly fragmentation; and **f**, paralog-assignment swaps. Points show the mean across 200 fixed-seed replicates and ribbons show the 2.5–97.5% empirical quantiles. The truth set consists of 274 high-confidence G1 and Grif1614 anchor shells and therefore measures robustness to the specified perturbations rather than universal genome-wide accuracy.

### Supplementary Figure 3 | Reviewer-driven tests of recovery, sampling, denominators and transcript support

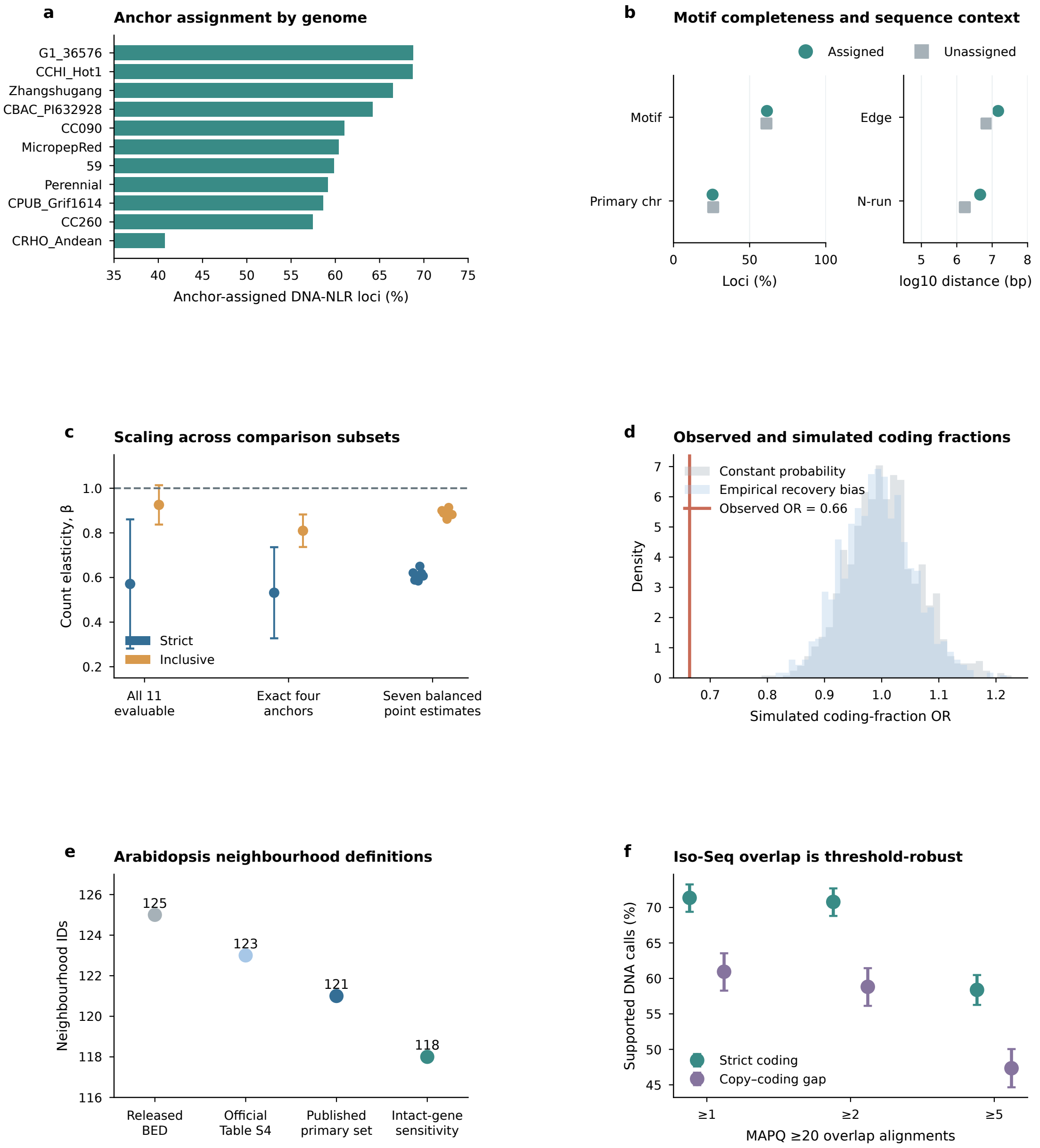


**a**, Fraction of all DNA-NLR loci assigned to anchor-defined neighbourhoods in each genome. **b**, Motif completeness, primary-sequence placement and distances from assembly edges and N-runs for assigned and unassigned loci; percentage and log10 distance measures are displayed on separate axes. **c**, Poisson log-link count elasticities in all-11-evaluable and exact-four-anchor cells, with two-sided neighbourhood-clustered 95% intervals, and all seven species-balanced point estimates. **d**, Null distributions of coding-fraction odds ratios under constant coding probability and the empirically measured copy-bin recovery profile; the magenta line is the observed odds ratio. **e**, Reconciliation of the released 125-neighbourhood BED, official 123-neighbourhood Table S4, reconstructed published 121-neighbourhood set and 118-neighbourhood intact-gene sensitivity subset. **f**, *Arabidopsis* DNA calls with at least one, two or five MAPQ ≥20 overlapping Iso-Seq alignments, with Wilson 95% intervals.

**Supplementary Figure 4 | Calibration and robustness of genome-wide copy-coding decoupling**

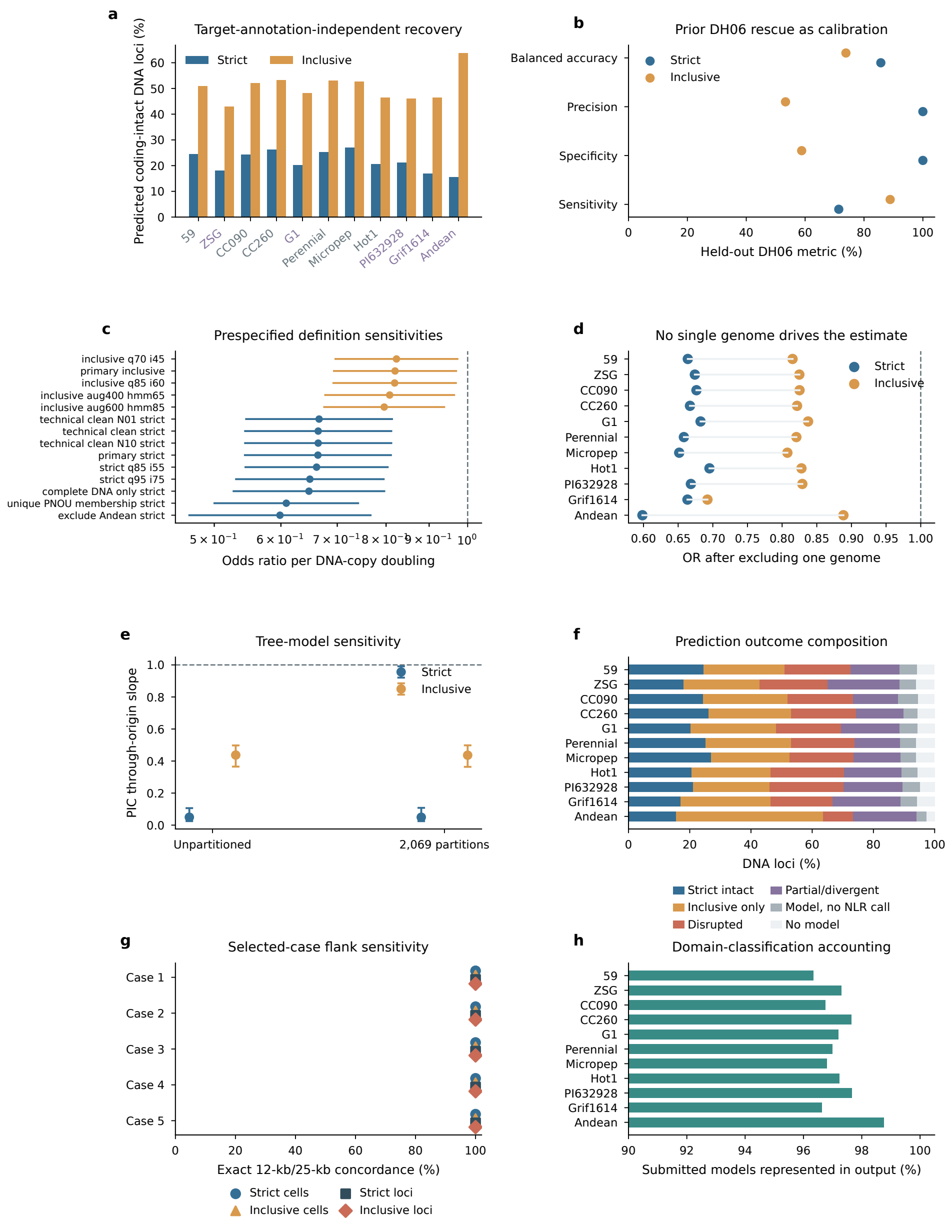


**a**, Per-genome strict and inclusive predicted coding-intact fractions from standardized leave-one-genome-out recovery. Purple labels identify genomes for which a matched annotation was available for calibration; those annotations were not used to call that genome. **b**, sensitivity, specificity, precision and balanced accuracy against the prior manually curated DH06 rescue set. **c**, odds ratios and 95% PNOU-clustered confidence intervals from every prespecified non-leave-one-genome-out binomial sensitivity model. **d**, primary odds-ratio estimates after excluding each genome in turn. **e**, through-origin PIC slopes and 95% PNOU-bootstrap confidence intervals under the unpartitioned and 2,069-partition species trees; the dashed line denotes equal numerical copy change, not a constant coding fraction. **f**, per-genome composition of strict intact, inclusive-only, explicitly disrupted, partial/divergent, model-without-NLR-call and no-model outcomes among DNA loci. **g**, exact locus- and cell-level concordance of strict and inclusive calls after rerunning all five selected cases with 25-kb rather than 12-kb flanks. **h**, fraction of submitted coding models represented in the domain-classification output. Unrepresented records are retained as no-calls and never recoded as positives.

**Supplementary Figure 5 | Phylogenetic turnover and repeat-source sensitivity across state-changing neighbourhoods**

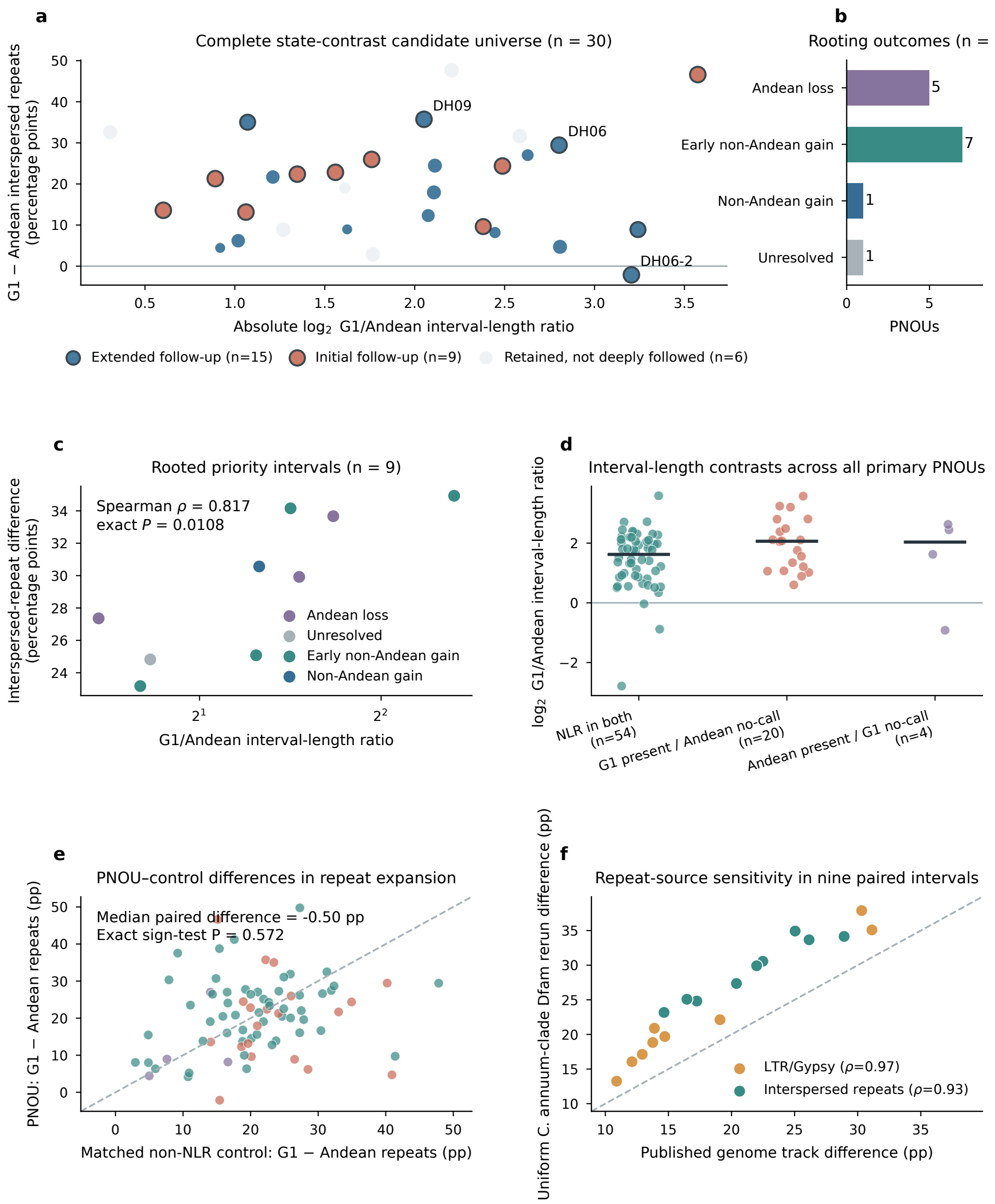


**a**, Complete 30-PNOU state-contrast universe; colour marks initial, extended or non-selected structural follow-up and dark outlines mark rooted cases. **b**, Rooted event classes among 14 directionally evaluated PNOUs. **c**, G1/Andean interval-length ratio and interspersed-repeat difference in nine initially prioritized rooted intervals. **d**, All observations and medians for three G1–Andean state classes among 78 primary PNOUs. **e**, Paired interspersed-repeat differences in PNOUs and length-matched four-anchor non-NLR controls. **f**, Comparison of published repeat tracks with a uniform *C. annuum*-clade Dfam 4.0 rerun in nine paired rooted intervals. These analyses position repeats as a broad structural background rather than the central driver of copy–coding–transcript divergence.

**Supplementary Figure 6 | Local small-variant context of the PI 585272 structural module**

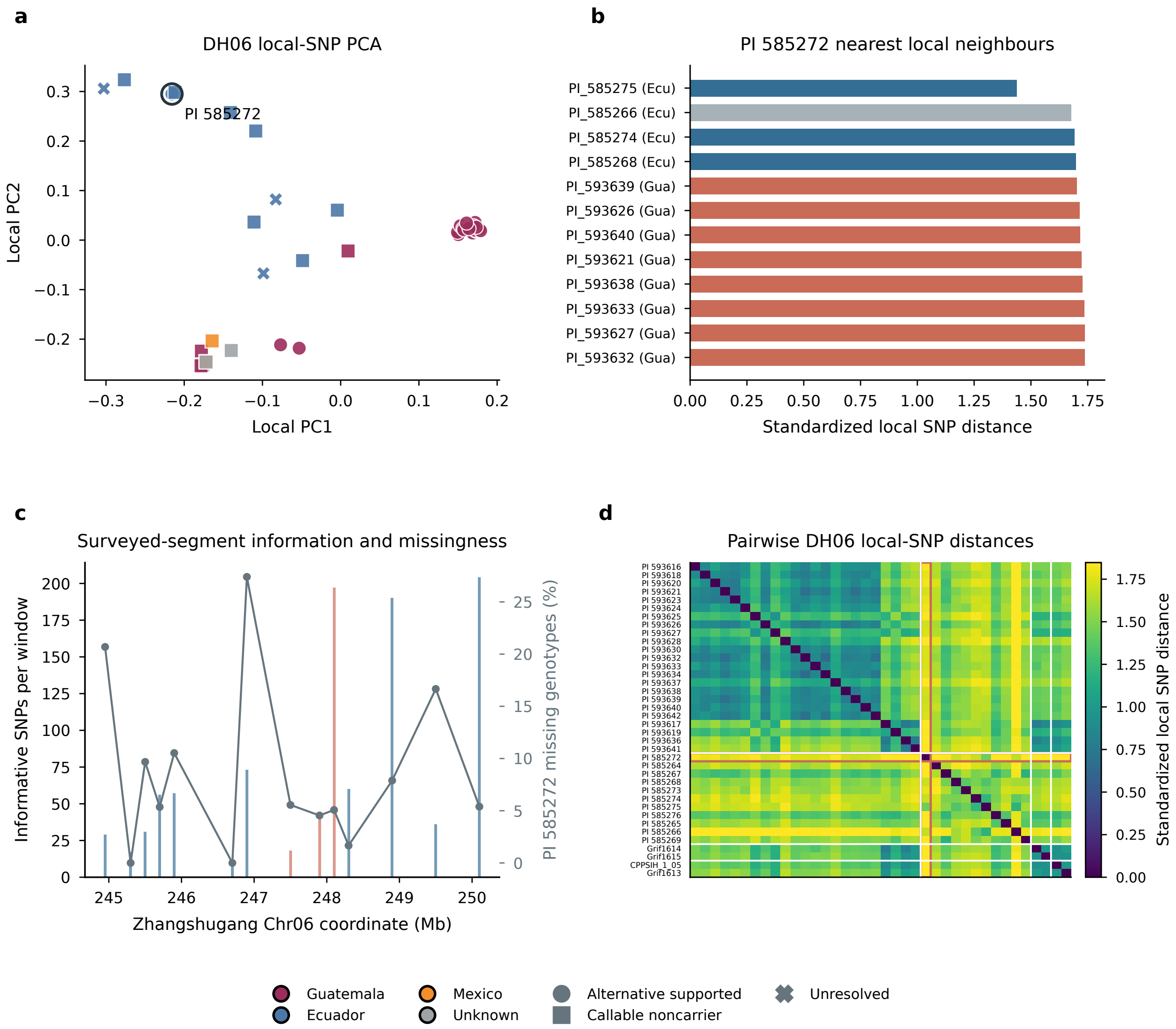


**a**, PCA of 185 physically thinned, DP/GQ-filtered common SNPs sampled across the DH06 interval and 1-Mb flanks; colour denotes origin, point shape denotes the revised three-state CROSS-COPY call and PI 585272 has a black outline. **b**, the 12 nearest accessions to PI 585272 by standardized local-SNP distance, coloured by the revised call state. **c**, informative-SNP count and PI 585272 missingness across the surveyed 50-kb segments; magenta segments overlap the independently projected cross-copy homology zone. **d**, pairwise local-SNP distance matrix ordered by origin and call state; the PI 585272 row and column are outlined. These exploratory local data show a mosaic background and are not used to infer a unique history of introgression, ancestral polymorphism or recurrent rearrangement.

**Supplementary Figure 7 | A callability-aware cross-copy structural module segregates within *C. pubescens***

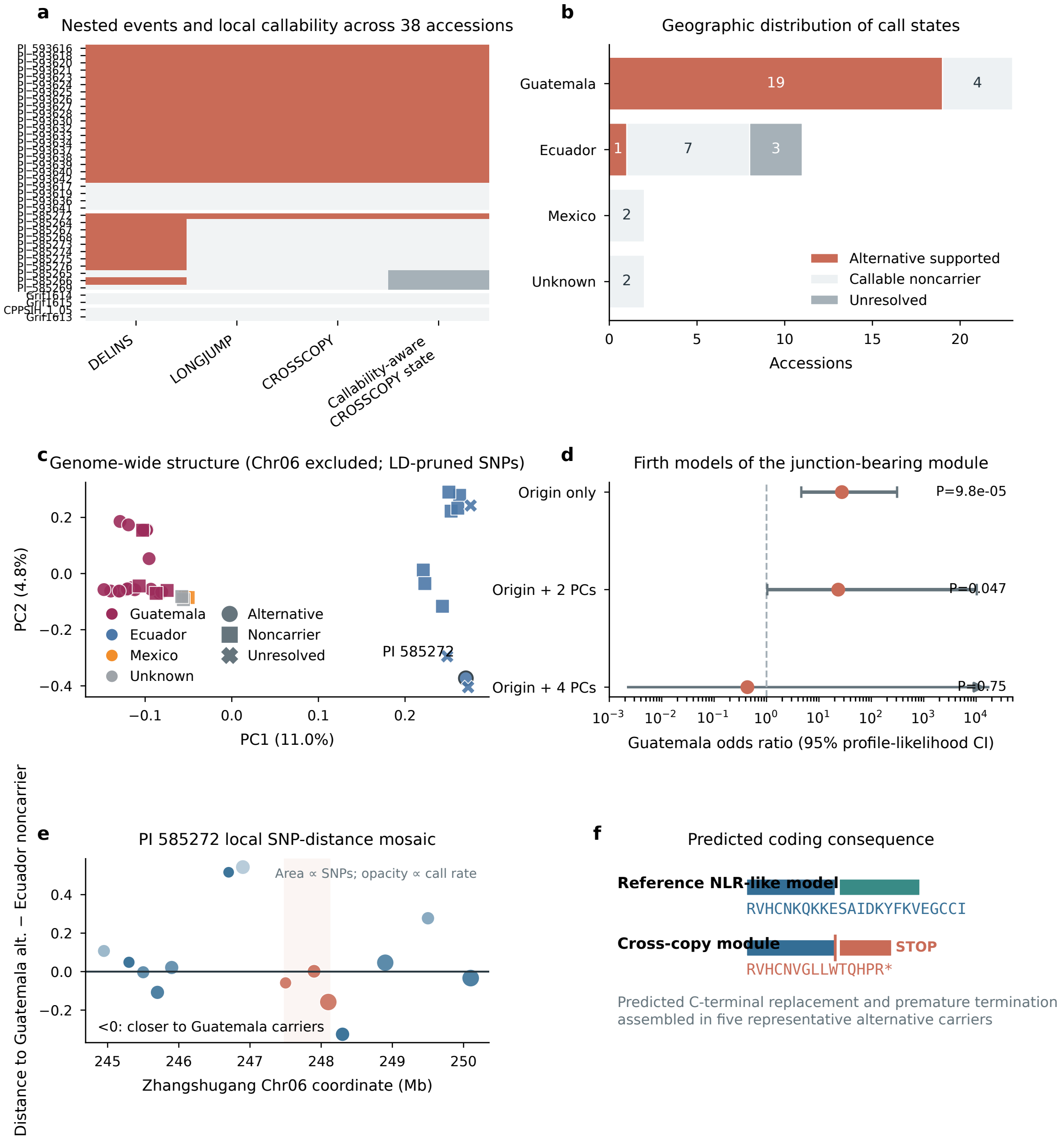


**a**, Nested junction evidence and three-state CROSSCOPY calls across 38 accessions. **b**, Origin-specific counts of alternative-supported, callable-noncarrier and unresolved samples; unknown origin is separate. **c**, PCA from Chr06-excluded, QC-filtered and LD-pruned genome-wide SNPs. Colour reports origin, shape reports call state and PI 585272 is outlined. **d**, Guatemala odds ratios and 95% penalized profile-likelihood confidence intervals from Firth models with zero, two or four PCs; the arrow for the four-PC model denotes a non-estimable upper profile limit. **e**, Exploratory contrast across surveyed 50-kb segments between the distance of PI 585272 to Guatemalan alternative carriers and callable Ecuadorian noncarriers. Point area reports informative SNP count, opacity reports PI 585272 call rate and the shaded interval overlaps the cross-copy homology zone. **f**, Junction-derived predicted terminal peptide replacement after the shared `RVHCN` sequence; the asterisk denotes premature termination. Population structure and geography are contextual analyses; the supported conclusion is segregation of the junction-defined structural module and its predicted coding consequence.

**Supplementary Figure 8 | Full PNOU length and repeat-sequence controls**

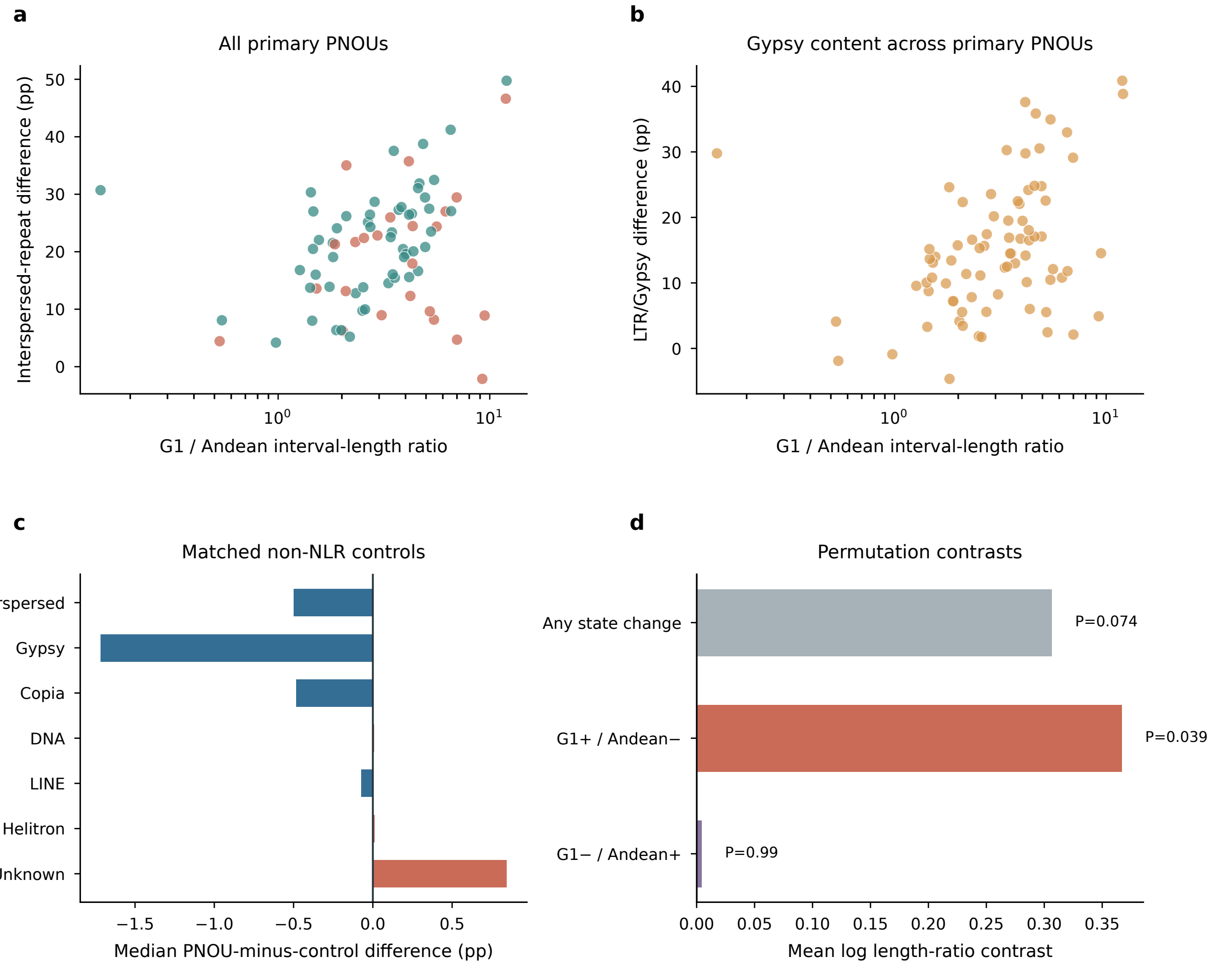


**a**, G1-to-Andean interval-length ratio and between-genome difference in total interspersed-repeat content for all 78 primary PNOUs. **b**, Equivalent comparison for LTR/Gypsy content. **c**, Median paired difference-in-differences between each PNOU and its unique length-matched, four-anchor, NLR-free control for seven repeat classes. **d**, Observed mean log-length-ratio contrasts and permutation *P* values for the three NLR-call comparisons. Percentage-point differences are denoted pp; non-significance is not interpreted as proof of equivalence.

**Supplementary Figure 9 | DH06 family assignment, annotation support and Andean continuity**

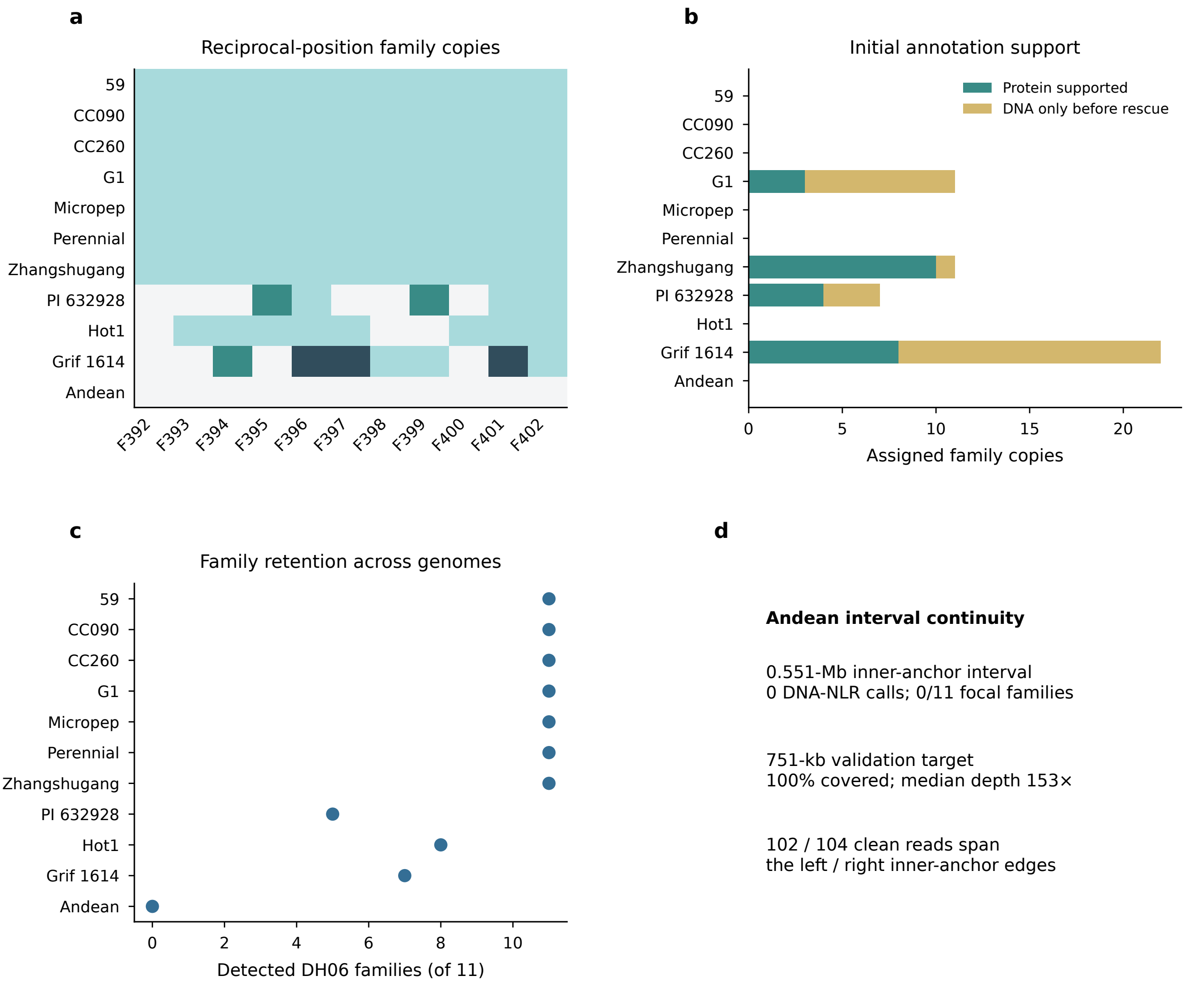


**a**, Copy-number heat map for reciprocal-position families F392–F402 across all 11 DH06 intervals. **b**, Protein-supported and DNA-only family-assigned copies before targeted coding-model analysis. **c**, Number of reciprocal-position families detected in each assembly. **d**, Independent raw-read evidence for the Andean interval: coverage of the validation interval, depth across 5-kb bins and primary reads spanning both inner-anchor boundaries. The Andean state is therefore an evaluable NLR no-call rather than an assembly-gap call.

**Supplementary Figure 10 | HiFi evidence for structural breakpoints and lesion sequence contexts**

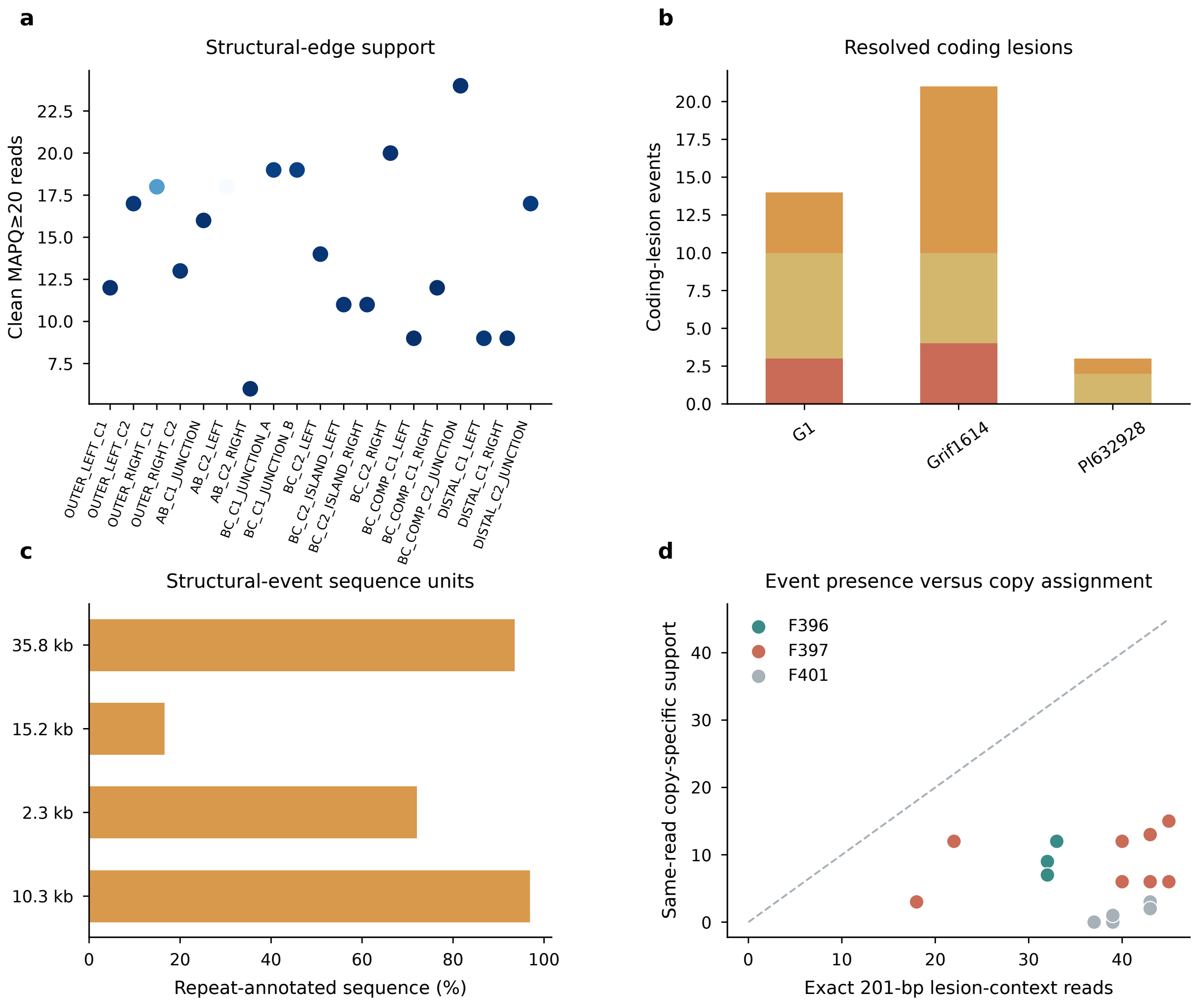


**a**, Clean MAPQ ≥20 PacBio reads spanning each Grif1614 structural edge. **b**, Frameshift and in-frame stop events among coding-disrupted DH06 loci. **c**, Repeat-annotated fractions of the four sequence-composition units representing five large structural events. **d**, Exact 101- and 201-bp lesion-centred context counts. These event-level counts establish lesion-sequence presence; physical-copy assignment is evaluated separately by same-read phasing in Fig. 5f and Supplementary Tables 25 and 44.

**Supplementary Figure 11 | Callability and genome-wide population-structure diagnostics**

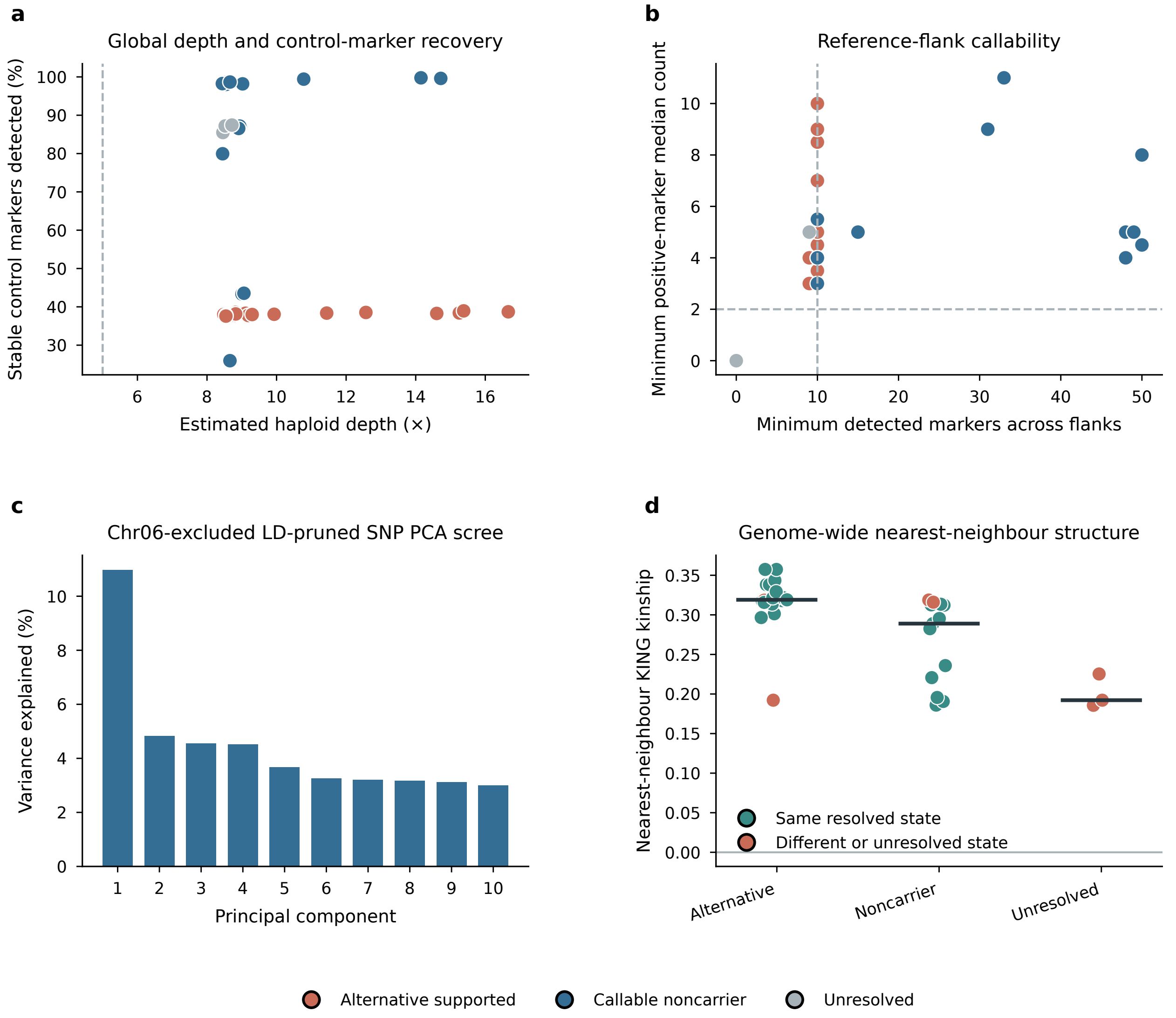


**a**, Estimated haploid depth and recovery of stable control markers for all 38 accessions. **b**, Minimum reference-marker support across the two CROSSCOPY flanks; dashed lines mark the prespecified noncarrier thresholds of ten detected markers and a positive-marker median count of two per flank. **c**, Variance explained by the first ten principal components from Chr06-excluded, QC-filtered and LD-pruned genome-wide SNPs. **d**, Nearest-neighbour KING kinship by local call state; colour records whether both resolved neighbours share the same CROSSCOPY state.

### Supplementary table titles and contents

**Supplementary Table 1 | Eleven structural assemblies and DNA-NLR catalogue**

Material identity, public accessions, source database, assembly checksum, assembly size, DNA-NLR counts, motif-completeness classes, sequence roles, N-run overlap and boundary-distance summaries.

**Supplementary Table 2 | Stable-anchor projection and order conservation**

Expected, uniquely placed, ambiguous and missing anchor HOGs; primary-placement rate and adjacent-anchor order consistency for every structural assembly.

**Supplementary Table 3 | Exact-ID concordance with the 2026 pepper pan-NLRome**

Exact-protein-ID intersections and reciprocal support fractions for PM1533, G1 and Zunla.

**Supplementary Table 4 | Complete 11-genome by 173-PNOU state matrix**

The unfiltered 11-genome × 173-PNOU matrix in phylogenetic tip order, retaining complete, partial, no-call and not-evaluable states.

**Supplementary Table 5 | Genome-wide locus coding predictions**

Target-annotation-held-out strict and inclusive calls, explicit disruption evidence, PNOU membership and technical flags for all 6,555 DNA-NLR loci.

**Supplementary Table 6 | PNOU-by-genome DNA and coding-copy matrix**

DNA, strict coding-intact and inclusive coding-intact counts for every positionally evaluable cell, retaining zero-copy cells and explicit non-evaluable states.

**Supplementary Table 7 | Global copy-coding summary**

Released-universe counts, gap frequencies, PNOU-bootstrap intervals and branch-event totals.

**Supplementary Table 8 | PNOU copy-coding summary**

Per-PNOU cell counts, DNA and coding-copy ranges, gap replication and phylogenetic variability.

**Supplementary Table 9 | Fixed-effect copy-coding models**

Primary, threshold, technical, membership and leave-one-genome-out binomial estimates with PNOU-clustered intervals with convergence and sample-size fields.

**Supplementary Table 10 | Log-link count elasticity and all sensitivity/influence fits**

Poisson log-link count elasticities with log DNA, neighbourhood/accession effects, clustered intervals and all sensitivity/influence fits. Legacy log1p coefficients are descriptive and remain in the source archive.

**Supplementary Table 11 | High-copy distribution and held-out recovery**

The complete cell-count distribution, locus recovery proportions and every adjusted bin contrast; the original locus records are separately retained as source data.

**Supplementary Table 12 | Pepper leave-one-neighbourhood influence**

All 522 complete-data/omission fits and three definition-level summaries for pepper neighbourhood influence.

**Supplementary Table 13 | Orthogonal high-confidence tests**

Complete-motif and complete/unique/clean subset definitions, fixed-effect estimates, leave-one-genome-out ranges, phylogenetic slopes, range summaries and DNA-gain classifications.

**Supplementary Table 14 | Complete-motif copy-coding matrix**

PNOU×genome DNA, strict and inclusive coding counts after retaining only complete-motif loci.

**Supplementary Table 15 | Complete/unique/clean copy-coding matrix**

The most restrictive PNOU×genome matrix after unique membership and N-fraction ≤0.01 filtering.

**Supplementary Table 16 | Frozen-screen candidate universe**

All cases passing or failing the prespecified DH06-excluded replication screen and the variables determining selection.

**Supplementary Table 17 | Five selected replication cases**

Frozen case identities, chromosomes, principal contrasts, branch-direction status and interpretation boundary.

**Supplementary Table 18 | Selected-case copy profiles**

Per-genome DNA, strict and inclusive coding-intact counts and evaluability for the five selected PNOUs.

**Supplementary Table 19 | Flank-width sensitivity**

Complete 12-kb versus 25-kb locus and cell comparisons, selected-case summaries and rerun accounting.

**Supplementary Table 20 | Evidence grade for non-DH06 replicated copy-coding cases**

Frozen DH01 and DH09 copy–coding architectures, raw-read and independent-reassembly availability, breakpoint status and manuscript claim boundaries.

**Supplementary Table 21 | Arabidopsis curated versus DNA count domains**

The complete Arabidopsis curated-versus-DNA count relation, retaining DNA-zero cells and the distinct count-model domains.

**Supplementary Table 22 | Complete state-contrast candidate universe and selection disclosure**

All 30 state-contrast PNOUs with selection wave, post hoc status, ranking variables, 11-genome occupancy, structural contrasts, rooted outcome and claim boundary.

**Supplementary Table 23 | Rooted PNOU events for all 14 directionally evaluated candidates**

Anchor signatures, outgroup states, parsimonious event class, confidence, interval ratios and repeat measurements for the 14 directionally evaluated candidates.

**Supplementary Table 24 | DH06 coordinates relative to the published Zhangshugang Chr06 introgression interval**

Zhangshugang coordinates and interval overlaps for the Liu *et al.* Chr06 introgression interval, project-local WP1C subregion, broad DH06 anchor shell, F392–F402 core, Grif1614 primary homology segments and CROSSCOPY breakpoint flanks.

**Supplementary Table 25 | Copy-specific HiFi phasing at F396, F397 and F401**

Same-read phasing of every shared lesion to the two physical F396, F397 and F401 loci, including strict alignment-score-margin support.

**Supplementary Table 26 | Damaged duplicate sequence context and lesion positions**

DH06 paired genomic-span/flank comparisons, lesion positions, base-level differences, extraction inputs and six-pair structural context.

**Supplementary Table 27 | Callability-aware CROSSCOPY genotypes with sample metadata**

Public sample metadata, global depth, stable controls, alternative-edge support, 50-marker reference-flank evidence and the three-state CROSSCOPY genotype for 38 accessions.

**Supplementary Table 28 | Public transcript accessions and selected cases**

Matched pepper RNA-seq runs, *Arabidopsis* Iso-Seq mapping QC and transcript support in five independent cases per species.

**Supplementary Table 29 | Released Arabidopsis input accounting and frozen definitions**

Frozen accession, DNA-call and curated-model denominators before reconciliation of the released 125-ID superset with the published 121-ID set.

**Supplementary Table 30 | Published-121 Arabidopsis copy-coding models**

Cluster-robust coding-fraction models and neighbourhood-clustered count elasticities in the reconstructed primary set.

**Supplementary Table 31 | Selected Arabidopsis replication cases**

Five independent neighbourhoods and their DNA, complete-motif and curated coding-copy profiles.

**Supplementary Table 32 | Matched pepper transcript summary and tests**

Frozen coding classes, strict/inclusive RNA-seq support and Fisher exact comparisons in G1, Andean and the combined panel.

**Supplementary Table 33 | Pepper locus-level transcript evidence**

DNA-NLR coordinates, frozen coding class and matched-tissue transcript-support calls.

**Supplementary Table 34 | Arabidopsis Iso-Seq support**

Accession-matched interval-overlap evidence for calls in the released 125-ID superset; the published-121 structure-level analysis is in Supplementary Table 41.

**Supplementary Table 35 | Held-out coding recovery by DNA-copy bin**

Recovery counts and Wilson intervals across DNA-copy bins, plus copy-number-adjusted models for 1,023 withheld complete gene models.

**Supplementary Table 36 | Anchor-assigned versus unassigned DNA-NLR loci**

Per-genome assignment fractions and assigned-versus-unassigned locus summaries, hypothesis tests and effect sizes.

**Supplementary Table 37 | Denominator-coupling null simulations**

Summary statistics and all simulated odds-ratio draws under constant coding probability and empirical copy-bin recovery bias.

**Supplementary Table 38 | Arabidopsis 125/123/121/118 neighbourhood reconciliation**

Count accounting, identifier-level membership, the four exclusions from the released BED superset and the three published-set neighbourhoods without a curated intact NLR gene.

**Supplementary Table 39 | Arabidopsis neighbourhood-set sensitivity models**

Coding-fraction and direct-count models in the reconstructed published 121-ID set, released 125-ID superset and 118-ID intact-gene subset.

**Supplementary Table 40 | Covariate-adjusted pepper and Arabidopsis transcript models**

Pepper locus and locus-by-tissue models and *Arabidopsis* Iso-Seq overlap models across read-count thresholds.

**Supplementary Table 41 | Arabidopsis locus-unique read and transcript-structure evidence**

Call-level and aggregate locus-unique read, splice, TAMA transcript and TransDecoder complete-ORF evidence, with covariate-adjusted models.

**Supplementary Table 42 | Methodological and coordinate positioning against recent NLR studies**

Explicit analytical and coordinate distinctions among Teasdale 2025, Dong 2026, Liu 2023 and CaNLOG.

**Supplementary Table 43 | Published-121 neighbourhood influence, sensitivity, range and cross-system concordance**

Leave-one-neighbourhood-out influence ranges and row-level refits, DNA-variable and top-five-excluded models, explicit curated-status definitions, range attenuation, and the descriptive two-system strict-fraction comparison.

**Supplementary Table 44 | Event-level copy-specific lesion support**

Exact 201-bp context counts and same-read physical-copy alignment evidence for each lesion entry.

**Supplementary Table 45 | P1 representative proteins and protein-NLR catalogue summary**

P1 representative-protein totals, raw protein classifier classes and explicit exclusions; the complete 5,083-record per-protein catalogue is supplied as tab-delimited source data.

**Supplementary Table 46 | Controlled-perturbation method definitions**

Positional frame, annotation dependency and assembly-break handling for the four compared representations.

**Supplementary Table 47 | Controlled-perturbation benchmark summary**

Mean recovery, precision, specificity, false-loss and unresolved fractions with 2.5–97.5% replicate quantiles.

**Supplementary Table 48 | All controlled-perturbation replicate metrics**

All 12,000 fixed-seed perturbation replicate × method results.

**Supplementary Table 49 | High-confidence anchor-shell benchmark units**

The 274 high-confidence G1 and Grif1614 anchor shells used as the conditional truth set.

**Supplementary Table 50 | Benchmark design boundaries and limitations**

Design boundaries that restrict interpretation of the controlled stress test.

**Supplementary Table 51 | Candidate-selection threshold sensitivity**

Candidate counts across 11-genome occupancy, interval-length and repeat-difference threshold combinations.

**Supplementary Table 52 | PNOU-level phylogenetic turnover classifications**

Evaluability, parsimony score and invariant, single-transition, repeated-transition or insufficient class for all 173 PNOUs.

**Supplementary Table 53 | Sample-level PNOU state summaries**

Complete, partial, no-call and not-evaluable burden in each structural genome.

**Supplementary Table 54 | Branch-level NLR-neighbourhood turnover burden**

Unambiguous gain, loss and ambiguous-change counts on every rooted species-tree branch.

**Supplementary Table 55 | All unambiguous and ambiguous optimal branch reconstructions**

PNOU-by-branch state-pair reconstructions, including ambiguous optimal histories rather than only selected events.

**Supplementary Table 56 | Repeat-source sensitivity in rooted intervals**

Per-locus differences from published genome tracks and a uniform *C. annuum*-clade Dfam 4.0 rerun.

**Supplementary Table 57 | Full primary G1-Andean PNOU interval set**

All 78 high- or medium-confidence G1–Andean intervals used for structural and repeat analyses.

**Supplementary Table 58 | Length-matched four-anchor non-NLR controls**

Unique length-matched, four-anchor, NLR-free controls and paired repeat contrasts.

**Supplementary Table 59 | DH06 interval evidence across 11 structural genomes**

Anchor intervals, DNA-NLR counts, family assignments, coding-model evidence and local-alignment summaries across 11 genomes.

**Supplementary Table 60 | DH06 reciprocal-position family copy matrix**

Material-by-family reciprocal-position copy counts for F392–F402.

**Supplementary Table 61 | Per-locus DH06 family and coding-model evidence**

Per-locus positional assignment, annotation support, rescue outcome, predicted coding state and local evidence.

**Supplementary Table 62 | Classification of DNA-only reciprocal-family loci**

Targeted classification outcomes for the 26 reciprocal-family loci lacking reliable initial models.

**Supplementary Table 63 | Coding-disrupted DH06 loci**

Coordinates, frameshift and in-frame stop counts, protein coverage and identity for resolved coding-disrupted loci.

**Supplementary Table 64 | Grif1614 duplicate family-pair evidence**

Physical-copy spacing, protein identity, local-tree relationship and reciprocal 100-kb alignment evidence.

**Supplementary Table 65 | Large structural differences between duplicated blocks**

Five large differences between the two Grif1614 blocks with exact coordinates and alignment provenance.

**Supplementary Table 66 | Raw-read structural-edge and lesion-context support**

Long-format read support for structural edges and lesion-centred sequence contexts.

**Supplementary Table 67 | Repeat composition of structural-event sequence units**

Repeat-union coverage, dominant class and family for four sequence-composition units.

**Supplementary Table 68 | Nested structural-event signatures in 38 accessions**

DELINS, LONGJUMP and CROSSCOPY signatures, descriptive H1–H5 junction-profile labels and focused-assembly support; H1–H5 are not independently phased haplotypes.

**Supplementary Table 69 | Firth models of Guatemala origin after SNP-structure adjustment**

Guatemala-origin odds ratios, confidence intervals and penalized likelihood-ratio tests with zero, two and four standardized PCs.

**Supplementary Table 70 | Chr06-excluded LD-pruned genome-wide SNP PCA**

Ten PCs, explained variance, origin and three-state CROSSCOPY call after excluding Chr06.

**Supplementary Table 71 | KING nearest-neighbour event concordance**

Genome-wide kinship nearest neighbour, origin, structural state and event concordance for every accession.

**Supplementary Table 72 | Genome-wide population-structure filters and variant counts**

Variant counts and the pre-specified DP, GQ, missingness, MAF, LD and chromosome-exclusion filters.

**Supplementary Table 73 | PI 585272 callability-aware local segment contrasts**

Window-level distance contrasts and marker completeness across DH06.

**Supplementary Table 74 | Predicted CROSSCOPY C-terminal coding consequence**

Breakpoint position, affected model, reference and alternative terminal sequences and evidence-calibrated coding prediction.

**Supplementary Table 75 | External DH06 gene-TE cross-check**

Exact comparison with the NLR-within-2-kb-TE table of Dong *et al.*; absence from that table is not treated as a de novo negative TE call.

**Supplementary Table 76 | Software and reference-resource versions**

Versions, analytical roles, local version evidence and citation keys for major programs and reference resources.

**Supplementary Table 77 | Uniqueness of reference-flank callability markers**

Whole-genome exact-hit counts and selection status for reference-flank markers.

**Supplementary Table 78 | NB-ARC sequence-cluster threshold sensitivity**

Recovery under 60%, 70% and 80% NB-ARC clustering thresholds.

**Supplementary Table 79 | Genotype-filter semantics and threshold pilot**

Retained variant counts under within-sample DP/GQ filters, a DP-only alternative, relaxed DP/GQ thresholds, no genotype filter and the rejected site-wide logical expression.

**Supplementary Table 80 | BUSCO species-tree topology sensitivity to locus partitioning**

Topology identity and branch support under unpartitioned and 2,069-partition BUSCO supermatrix models, together with the number of PNOU classifications affected.

**Supplementary Table 81 | DH06 local-SNP PCA with population metadata**

Local principal-component coordinates, origin, revised three-state call and resolved comparator-group status for all 38 accessions.

**Supplementary Table 82 | PI 585272 nearest neighbours from DH06 local SNPs**

Ranked standardized local-SNP distances from PI 585272 to the remaining accessions.

**Supplementary Table 83 | Analytically checkable validation of the Firth implementation**

Agreement with the analytically checkable Haldane–Anscombe/Firth odds ratio in a separation-like 2×2 table, agreement with direct penalized-likelihood optimization and profile-likelihood interval convergence.

**Supplementary Table 84 | Range-attenuation tests**

Exact one-sided sign tests comparing DNA-copy range with strict and inclusive coding-intact-copy range across all DNA-variable PNOUs.

**Supplementary Table 85 | Phylogenetic independent contrasts**

All branch contrasts together with strict and inclusive through-origin estimates and PNOU-cluster bootstrap intervals.

**Supplementary Table 86 | DNA-gain event classes**

Strict and inclusive counts and proportions of proportional, partially decoupled, fully decoupled and mixed/uncertain events among unambiguous DNA-gain branches.

**Supplementary Table 87 | Joint branch reconstructions**

PNOU-by-branch DNA and coding state pairs from the lexicographic stepwise-Sankoff reconstruction, retaining alternative optimal histories.

**Supplementary Table 88 | Held-out DH06 calibration**

Locus-level truth comparisons and aggregate sensitivity, specificity, precision, accuracy and balanced accuracy for both coding definitions.

**Supplementary Table 89 | Sample-level coding-recovery QC**

Prediction outcome fractions, matched-annotation availability and domain-classification accounting for each genome.

**Supplementary Table 90 | Scientific consistency checks**

Machine-readable invariant checks for non-negative counts, coding≤DNA constraints, denominator consistency and exact selected-case flank concordance.

### Evidence and interpretation boundaries

The complete data release distinguishes direct sequence observations, model-derived coding predictions, locus-level transcript support and evolutionary reconstructions. External call-set overlap is not labelled validation; lesion-context presence is not equated with physical-copy phasing; an absent alternative junction is not a noncarrier without reference-flank callability; and unknown geographic origin remains a separate class. The controlled benchmark evaluates specified failure modes within a high-confidence anchor-shell set, while the repeat rerun evaluates source sensitivity. Regression-level sublinear scaling is concordant in two independently defined systems, whereas *Arabidopsis* neighbourhood-level range attenuation is not significant and no plant-wide meta-analysis is claimed. DH06 is coordinate-distinct from the published Zhangshugang Chr06 introgression core, and non-DH06 selected cases remain replicated copy–coding patterns unless independently resolved. Strict and inclusive predicted coding-intact states are sequence-based bounds, and RNA-seq/Iso-Seq support does not establish functional receptor activity. Resistance phenotype and local adaptation remain experimental questions.
